# Using a GPT-5-driven autonomous lab to optimize the cost and titer of cell-free protein synthesis

**DOI:** 10.64898/2026.02.05.703998

**Authors:** Alexus A. Smith, Edmund L. Wong, Ronan C. Donovan, Brad A. Chapman, Ryan Harry, Pooyan Tirandazi, Paulina Kanigowska, Elizabeth A. Gendreau, Robert H. Dahl, Michal Jastrzebski, Jose E. Cortez, Christopher J. Bremner, José C. Morales Hemuda, James Dooner, Ian Graves, Rahul Karandikar, Christopher Lionetti, Kevin Christopher, Andrew L. Consiglio, Alyssa Tran, William McCusker, Duy X. Nguyen, Isis Botelho Nunes da Silva, Alvaro R. Bautista-Ayala, Monica P. McNerney, Sean Atkins, Michael McDuffie, Will Serber, Bradley P. Barber, Trinh Thanongsinh, Andrew Nesson, Bibek Lama, Brandon Nichols, Cameron LaFrance, Tenzing Nyima, Alicia Byrn, Rashard Thornhill, Bryan Cai, Lizvette Ayala-Valdez, Alycia Wong, Austin J. Che, Walter Thavarajah, Daniel Smith, Thomas F. Knight, David W. Borhani, Jerry Tworek, Mostafa Rohaninejad, Ahmed El-Kishky, Nathan C. Tedford, Tejal Patwardhan, Yunxin Joy Jiao, Reshma P. Shetty

## Abstract

We used an autonomous lab, comprising a large language model (LLM) and a fully automated cloud laboratory, to optimize the cost efficiency of cell-free protein synthesis (CFPS). By conducting iterative optimization, the LLM-driven autonomous lab was able to achieve a 40% reduction in the specific cost ($/g protein) of CFPS relative to the state of the art (SOTA). This cost reduction was accompanied by a 27% increase in protein production titer (g/L). Iterative experimental design, experiment execution, data capture and analysis, data interpretation, and new hypothesis generation were all handled by the LLM-driven autonomous lab. The interface between OpenAI’s GPT-5 LLM and Ginkgo Bioworks’ cloud laboratory incorporated built-in validation checks via a Pydantic schema to ensure that AI-designed experiments were properly specified. Experimental designs were translated into programmatic specification of multi-instrument biological workflows by Ginkgo’s Catalyst software and executed on Ginkgo’s Reconfigurable Automation Cart (RAC) laboratory automation platform, with human intervention largely limited to reagent and consumables preparation, loading and unloading. By integrating LLMs with programmatic control of a cloud lab, we demonstrate that an LLM-driven autonomous lab can successfully perform a real-world scientific task, highlighting the potential of AI-driven autonomous labs for scientific advancement.

## 1. Introduction

There are two central bottlenecks in scientific research. The first is so-called “wet lab” experimentation: the hands-on, physical manipulation of chemical and biological materials to conduct experiments. The second is intelligence: the conception, planning, and design of experiments, and the analysis and interpretation of resulting data, to answer a given scientific question. Advances in laboratory automation, software architectures, application programming interfaces, and cloud labs promise to alleviate the former bottleneck. Similarly, rapid advances in large language models (LLMs), particularly reasoning models that can solve complex, multi-step problems by breaking them down into logical, sequential steps, can automate the latter. Together, reasoning models and “autonomous labs” are capable of performing all steps of routine scientific discovery: experimental planning, design, execution, data analysis and interpretation, and new hypothesis generation. Early efforts have illustrated the potential of autonomous labs across multiple disciplines including chemical research,^1^ material science^2–6^ and biological design.^7,8^ Until recently, reasoning LLMs have been deployed mostly in computationally verifiable tasks such as algorithms, math, and physics. Evaluating AI performance in designing biological experiments is particularly challenging due to the need for wet lab validation.^9^ As a result, the potential of AI-driven autonomous labs for advancing real-world scientific applications remains largely untapped.

One such real-world scientific application is cell-free protein synthesis (CFPS), a method for recombinant protein production of both laboratory and commercial importance that has proven difficult to optimize.^10–12^ Given the widespread use of CFPS in rapid prototyping^13^ including for antibodies,^14–16^ peptides,^17^ proteins,^18^ enzymes,^19^ and proteins for structure determination,^20^ as well as in the commercial production of an antibody-drug conjugate luveltamab tazevibulin (luvelta),^21,22^ improvements in CFPS achieved by the application of autonomous lab experimentation would demonstrate both the power of the autonomous lab concept and deliver real-world benefits.

The very large CFPS parameter space suggests that its optimization would be a suitable—and challenging—testbed for autonomous lab experimentation. CFPS reactions consist of many components (DNA template encoding the protein of interest, cell lysate, nucleic acids, amino acids, energy sources, cofactors and coenzymes, buffers, salts, etc.). CFPS can also produce proteins bearing a wide variety of modifications, including disulfide bonds,^23,24^ glycosylation,^25,26^ phosphorylation^27^ and non-canonical amino acids,^28–32^ further broadening the search space. Accordingly, there have been significant prior efforts leveraging machine learning and similar techniques to optimize CFPS reactions against two common figures of merit: titer (gram of protein produced per liter of reaction) and specific cost (the total reaction composition cost per gram of protein produced).^33–40^ Reducing specific cost—a critical factor given the much larger scale of experimentation made possible by autonomous labs—without sacrificing titer makes CFPS optimization a non-trivial problem.

Here we combined OpenAI’s GPT-5^41^ with Ginkgo Bioworks’ Reconfigurable Automation Cart (RAC)-based cloud laboratory in Boston, MA to perform autonomous lab experimentation with the objective of minimizing *E. coli* CFPS specific cost. The LLM-driven autonomous lab was able to produce a standard benchmark protein, superfolder green fluorescent protein (sfGFP), at a cost of $422/g versus a recent state of the art (SOTA) report of $698/g by Olsen *et al*. (2025),^42^ a 40% cost reduction over SOTA. This cost reduction was accompanied by an *increase* of 27% in protein production (titer, g/L) versus SOTA, confirming that cost and titer can be improved simultaneously without sacrificing one for the other.

## 2. Results

### We used a GPT-5-driven autonomous lab to optimize CFPS

Our approach to GPT5-driven autonomous lab experimentation, applied here to CFPS optimization, is illustrated in Figure 1. We used GPT-5^41^ to design *E. coli* lysate-based CFPS reaction compositions that optimize production of a test protein, superfolder green fluorescent protein (sfGFP), with the objective of minimizing specific cost. The first iteration of CFPS reactions (what we refer to in this work as *Step 0*) were designed by GPT-5 without the benefit of any example reactions or prior experimental results (a setting referred to as “zero-shot”); thus, GPT-5 only leveraged intrinsic knowledge stored in model weights to generate experimental reaction compositions. GPT-5-designed experiments were constrained to be properly specified, e.g. verified to be scientifically valid and physically executable on Ginkgo’s cloud lab, by applying a Pydantic-based validation schema.

**Figure 1.**
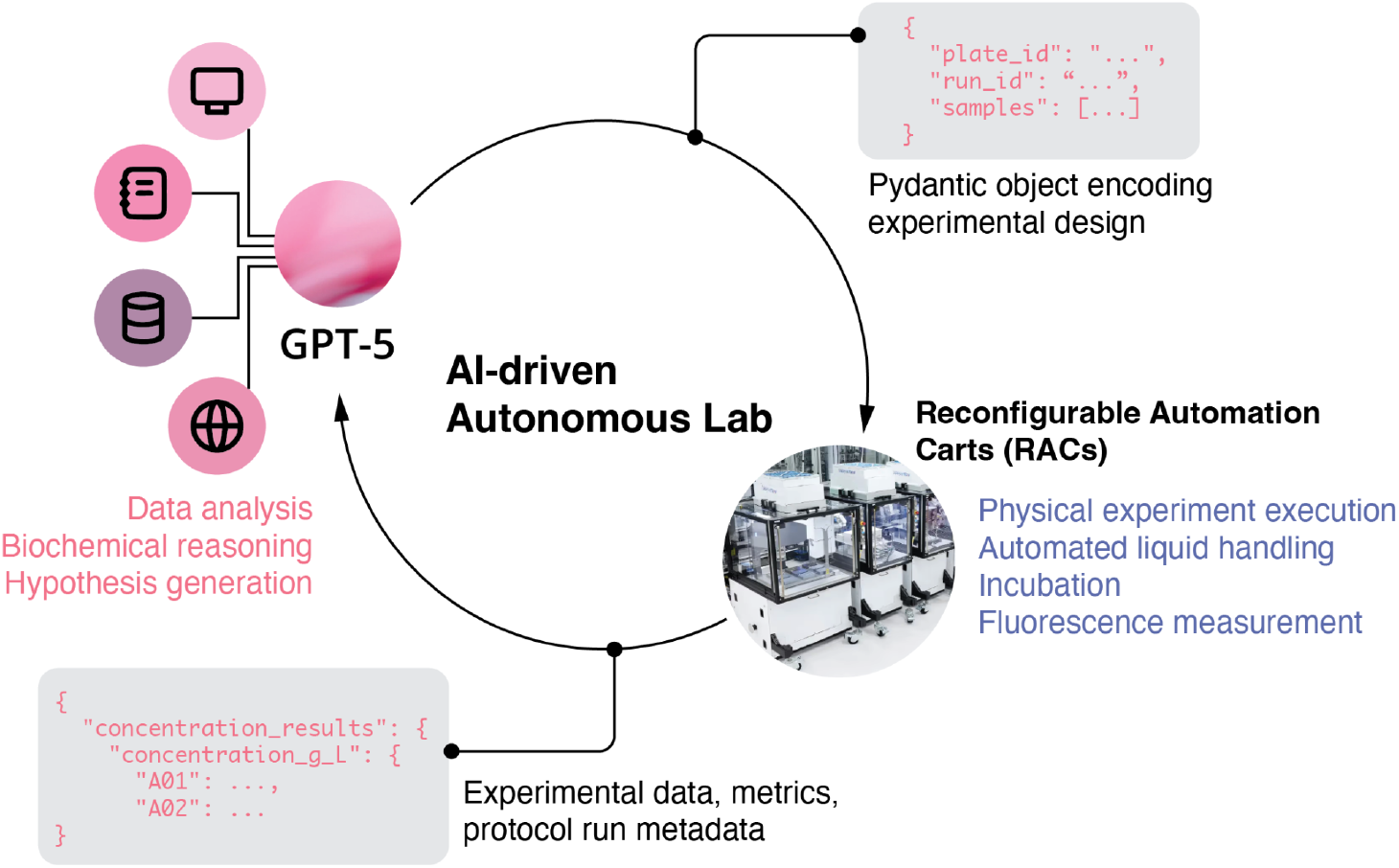
Experimental iterations by a GPT-5-driven autonomous lab implemented on RACs. OpenAI’s GPT-5 proposes a batch of experimental designs in 384-well plate format and validates them using a Pydantic schema. Designs are then programmatically converted to Catalyst protocol runs for execution on Ginkgo’s RAC-based cloud laboratory. After experiments are executed, (meta)data and metrics are returned to GPT-5 for analysis, interpretation, hypothesis generation, and design of the next round of experiments.

All CFPS reactions were run on Ginkgo’s Reconfigurable Automation Carts (RAC)-based cloud laboratory in Boston, MA. RACs are a modular laboratory automation solution developed by Ginkgo. Each RAC contains a single instrument type—e.g., liquid handler, centrifuge, incubator, plate reader—and standardized hardware that enables a RAC to move samples in SBS-format microtiter plates to and from that instrument (via a 6-axis robotic arm) and between RACs (via a sample transport track). RACs flexibly combine into unified assemblies to perform fully automated, scalable workflows. Ginkgo’s RAC command-and-control software, *Catalyst*, enables programmatic specification of multi-instrument workflows and orchestrates the running of many such workflows simultaneously on the same RAC assembly, including RAC and instrument unit operations, sample transport, and (meta)data collection (Supplemental Figure 1). Ginkgo’s cloud lab includes all the instrumentation needed for the described CFPS workflow (Supplemental Figure 2).

**Figure 2.**
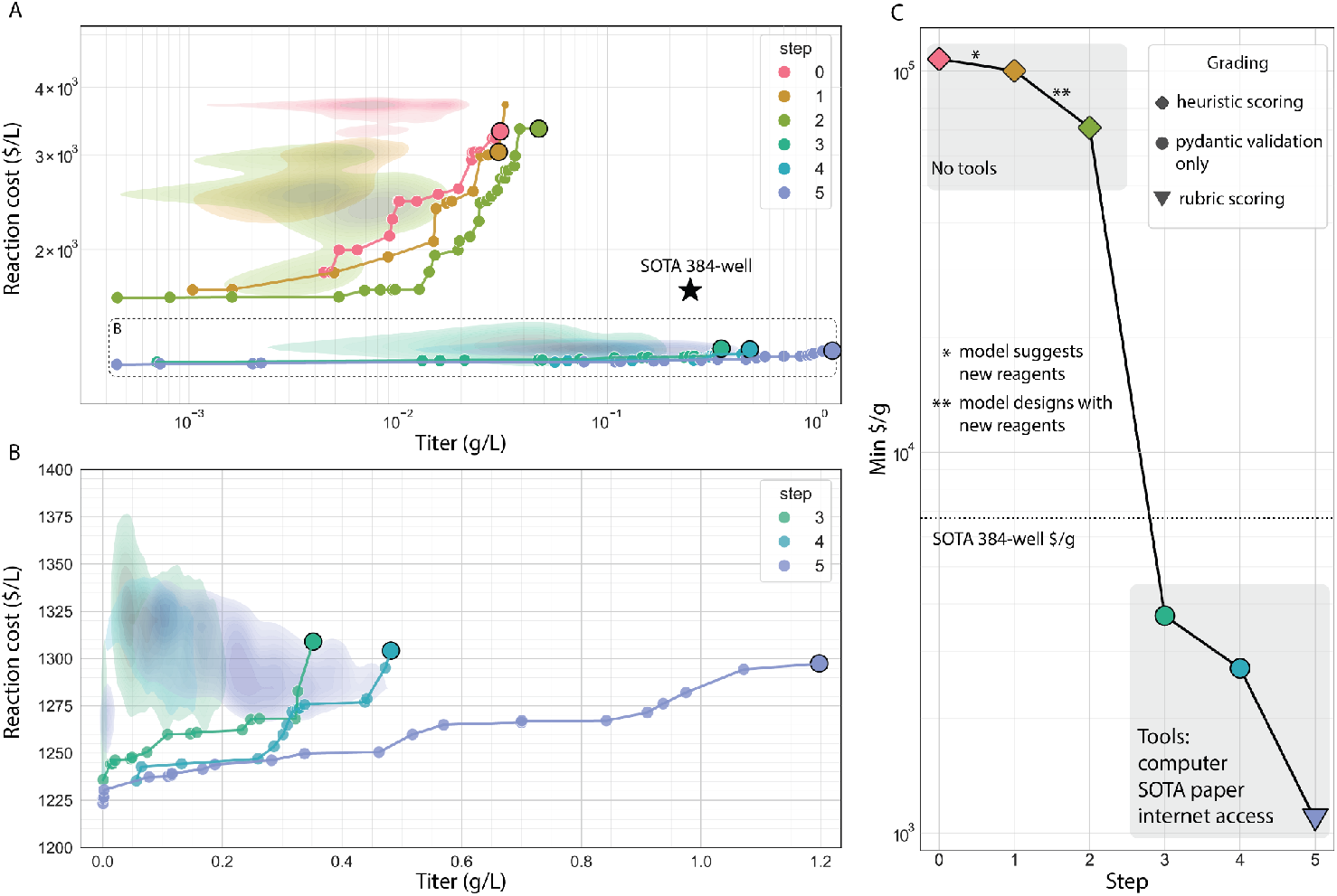
Autonomous lab experimentation reduces CFPS reaction cost and increases protein titer over six iterative steps. (A) CFPS reaction cost ($/L) versus protein titer (g/L) across *Steps 0-5* of experimentation in 384-well plate reaction geometry (Pareto frontier plotted on a log-log scale). Reactions representing the minimum specific cost ($/g) for each step are highlighted with larger dots. Star: SOTA in 384-well plates as reported by Olsen *et al*. (2025).^*42*^ (B) *Steps 3-5* of panel A replotted on a linear scale. (C) Best specific cost ($/g) obtained in 384-well plate reaction geometry at each step. Dotted line: SOTA in 384-well plates as reported by Olsen *et al*.

We ran CFPS reactions at 20-µL scale in 384-well plates. To improve dispensing accuracy and precision, different cloud lab liquid dispensing instruments were used to add reaction components (base buffer, reagents, DNA template and cell lysate) based on dispense volume and component viscosity (see Materials and Methods). Each plate consisted of 78 distinct reaction compositions (4 replicates each), purified sfGFP for absolute protein quantification, and positive and negative controls. The cloud lab performed reaction component addition, CFPS reaction incubation, and fluorescence data acquisition. We used custom software code to (a) translate GPT-5-designed experiments into Catalyst protocol runs (Supplemental Figure 3) on Ginkgo’s cloud lab and (b) retrieve and return data to GPT-5. Experimental data was returned to GPT-5 via a JSON file comprising protein titers. Human intervention in laboratory experiments was primarily limited to preparation, loading and unloading of reagents and consumables on and off the RAC system.

A compelling capability of an LLM-driven autonomous lab is its ability to generate human-readable laboratory notebook entries (Supplementary Material section 7.3). GPT-5 documented how it analyzed data, observations it made of experimental trends, surprising or confusing results, and new hypotheses it formulated for testing. Early entries from *Step 0*, for example, critiqued titer variability between replicates within a plate (CVs >40%). To reduce titer CVs, we modified certain reagent stocks to improve the precision and accuracy of liquid dispensing steps (Supplementary Material section 7.4). For example, we reduced concentrations to enable more accurate liquid transfer on the Echo 525 acoustic dispenser (potassium glutamate, magnesium glutamate, DMSO) and changed stock solutions to reduce precipitation (e.g., switched to a maltodextrin with a shorter chain length, and increased the pH of the tyrosine stock solution to pH 12). Ginkgo personnel performed quality testing of precipitation and Echo liquid handling of different reagent stock solutions manually. With these improvements, we were able to achieve median per-plate CVs of 17%, with 90% of initially screened plates having CVs less than 30%. In total in *Step 0*, 146 384-well plates totalling 7,502 unique reaction compositions were designed, tested and data returned to GPT-5.

In *Step 1*, GPT-5 used experimental data from *Step 0* to inform the next round of reaction designs. Thirty-two 384-well plates (1,656 unique reaction compositions) were executed on Ginkgo’s cloud lab with data returned to GPT-5. In this step, GPT-5 was able to design reaction compositions with reduced cost but similar titers versus *Step 0* (Figure 2A). We also noted that in many lab notebook entries, GPT-5 suggested additional reagents not already available in the reagent list that it reasoned might improve specific cost. Indeed, there were so many lab notebook entries, and GPT-5 had so many reagent suggestions, that we asked GPT-5 to analyze and rank-order its reagent suggestions (Supplementary Material section 7.5), sourced those reagents that were commercially available, and added them to the list of reagents available to the model for *Step 2* or *Step 3* (Supplementary Table 1).

In *Step 2*, although GPT-5 was able to make use of the additional reagents, it chose to do so relatively sparingly. Nevertheless, the GPT-5-driven autonomous lab achieved both lower reaction costs and higher titers versus *Step 0–1* (Figure 2A). In total in Step 2, 64 384-well plates corresponding to 3,586 unique reaction compositions were tested with data return to GPT-5. After *Step 2*, we made a series of process improvements to our CFPS workflow to improve CFPS reaction productivity. In particular, we switched to a different sfGFP plasmid design and improved our cell-free lysate preparation process (Supplementary Material section 7.4 and Materials and Methods).

In *Step 3*, we significantly expanded both the tools the model could access and the experimental (meta)data that we provided GPT-5. In particular, we provided GPT-5 with access to the internet, a computer, and data analysis packages. We also provided the model with the preprint on SOTA published by Olsen *et al*. (2025)^42^ on August 1, 2025 (including their supplementary materials). The model was able to access relevant scientific literature online. Finally, we exposed the model to additional, potentially useful experimental (meta)data, including raw fluorescence values, titers from positive and negative controls, reagent lot numbers, liquid handling errors, quality-of-fit metrics for per-plate standard curves, instrument protocols and parameters, and actual CFPS incubation times (which may vary from the intended incubation time of 20 h during periods of high cloud lab utilization). Together, these changes, along with improvements to the DNA template and cell lysate, enabled a jump in performance in both reaction cost and titer (Figure 2A), with the model making more use of the additional reagents. Note that we opted to forego heuristic scoring in this step. Top-performing reaction conditions from *Step 3* were k-means clustered to select a diversity of reagent compositions, down-selected by the $/g metric, and re-tested in 96-well plates (cloud lab) and 2-mL flat-bottom tubes (manually). As Olsen *et al*. note, CFPS reaction titers, and thus specific costs, vary with reaction oxygenation.^42^ Batch 20 µL CFPS reactions in a 2-mL tube format generally yielded higher titers than those in 96-well plates, which in turn generally yielded titers higher than those in 384-well plates, likely due to differences in oxygenation and known surface area to volume effects.^43^ Results from 96-well plates were provided to the model. In total, 48 384-well plates corresponding to 3,211 unique reaction compositions were tested in *Step 3*.

*Step 4* was performed by the GPT-5-driven autonomous lab using a similar approach as *Step 3* with 62 384-well plates being tested corresponding to 4,244 unique reaction compositions. GPT-5 was able to achieve lower reaction costs and higher protein titers in *Step 4* versus *Step 3* (Figure 2B). In *Step 5*, we adopted a rubric scoring approach and ran 128 384-well plates comprising 7,809 unique reaction compositions. Step 5 resulted in similar reaction costs but a jump in titers (Figure 2B). In total over six steps, we executed 480 384-well plate experimental designs comprising 29,527 unique reaction compositions in six months via the GPT-5 driven autonomous lab, with each successive step resulting in lower specific cost (Figure 2C).

### The GPT-5-driven autonomous lab improved CFPS specific cost by 40% versus SOTA

We benchmarked the performance of CFPS reaction compositions from the autonomous lab against the best-performing composition “RF_opt_” from Olsen *et al*. (2025),^42^ the current SOTA. Olsen *et al*. sought to reduce reagent costs–the cost of all CFPS reaction components other than cell lysate and DNA template. In light of Olsen *et al*.’s substantial reduction in reagent costs, however, cell lysate and DNA template now account for >90% of total CFPS reaction cost in SOTA. Accordingly, we focused on optimizing total CFPS reaction cost (specific cost, $ per gram of protein produced), i.e., the sum of reagent, lysate, and DNA costs. Indeed, we were able to further reduce reagent, lysate and DNA costs. Olsen *et al*. reported a total reaction cost of $1,676/L_CFE_ (their Supplemental Table 1) and a titer of 2.4 ± 0.3 g/L (their Table 1), resulting in a calculated specific cost of $698/g for RF_opt_. Note that we use the same cell lysate, DNA template and reagent unit costs as Olsen *et al*. for direct comparison of specific costs obtained here to SOTA (see Olsen *et al*. Supplemental Tables 1 and 2). If a unit cost for a particular reagent was not available from Olsen *et al*., we adopted their approach and determined the unit cost based on the largest retail unit available from the vendor.

**Table 1.** Top-performing CFPS reaction compositions identified in this work. All CFPS reactions were performed at the 20-µL scale in the designated reaction geometry (2-mL tubes, 96-well plate or 384-well plate). Reported errors are standard deviations of 8 replicates for 384-well plates and 4 replicates for 96-well and 2-mL tubes. Values for RF_opt_ (SOTA) are calculated from Olsen *et al*. (2025)^*42*^ Supplemental Table 1 (total reaction cost calculation of $1,676 per liter of CFPS reaction), their Table 1 (titer for 15-µL scale in 2-mL tubes), and their Supplemental Figure 7a (estimated titer for 15-µL scale in 384-well plates). Reaction composition 1777863_77 has the lowest specific cost in 2-mL tubes, with a 40% improvement over SOTA; however, reaction composition 1793122_35 demonstrates the most balanced performance across reaction geometries, with a 38% improvement over SOTA in 2-mL tubes and a 188% improvement over SOTA in 384-well plates.

| Reaction composition identifier | Specific Cost (\$/g) | Protein Titer (g/L) | | |
| --- | --- | --- | --- | --- |
|  | 2-mL tubes | 2-mL tubes | 96-well plates | 384-well plates |
| $RF_{opt}^*$ (SOTA) | \$698* | $2.39 \pm 0.28^*$ | Not applicable* | $\sim 0.25^*$ |
| 1777863_77 | \$422 | $3.04 \pm 0.30$ | $0.66 \pm 0.12$ | $0.29 \pm 0.01$ |
| 1784943_26 | \$425 | $3.01 \pm 0.27$ | $0.86 \pm 0.24$ | $0.33 \pm 0.05$ |
| 1793122_35 | \$430 | $3.04 \pm 0.18$ | $1.52 \pm 0.32$ | $0.72 \pm 0.38$ |
| 1793665_74 | \$446 | $2.92 \pm 0.19$ | $1.31 \pm 0.33$ | $0.49 \pm 0.05$ |
| 1793119_36 | \$466 | $2.77 \pm 0.19$ | $1.28 \pm 0.13$ | $0.78 \pm 0.58$ |
| 1794089_36 | \$470 | $2.75 \pm 0.27$ | $2.00 \pm 0.49$ | $0.53 \pm 0.07$ |

\* As reported by Olsen *et al.* (2025).<sup>42</sup>

**Table 2.**
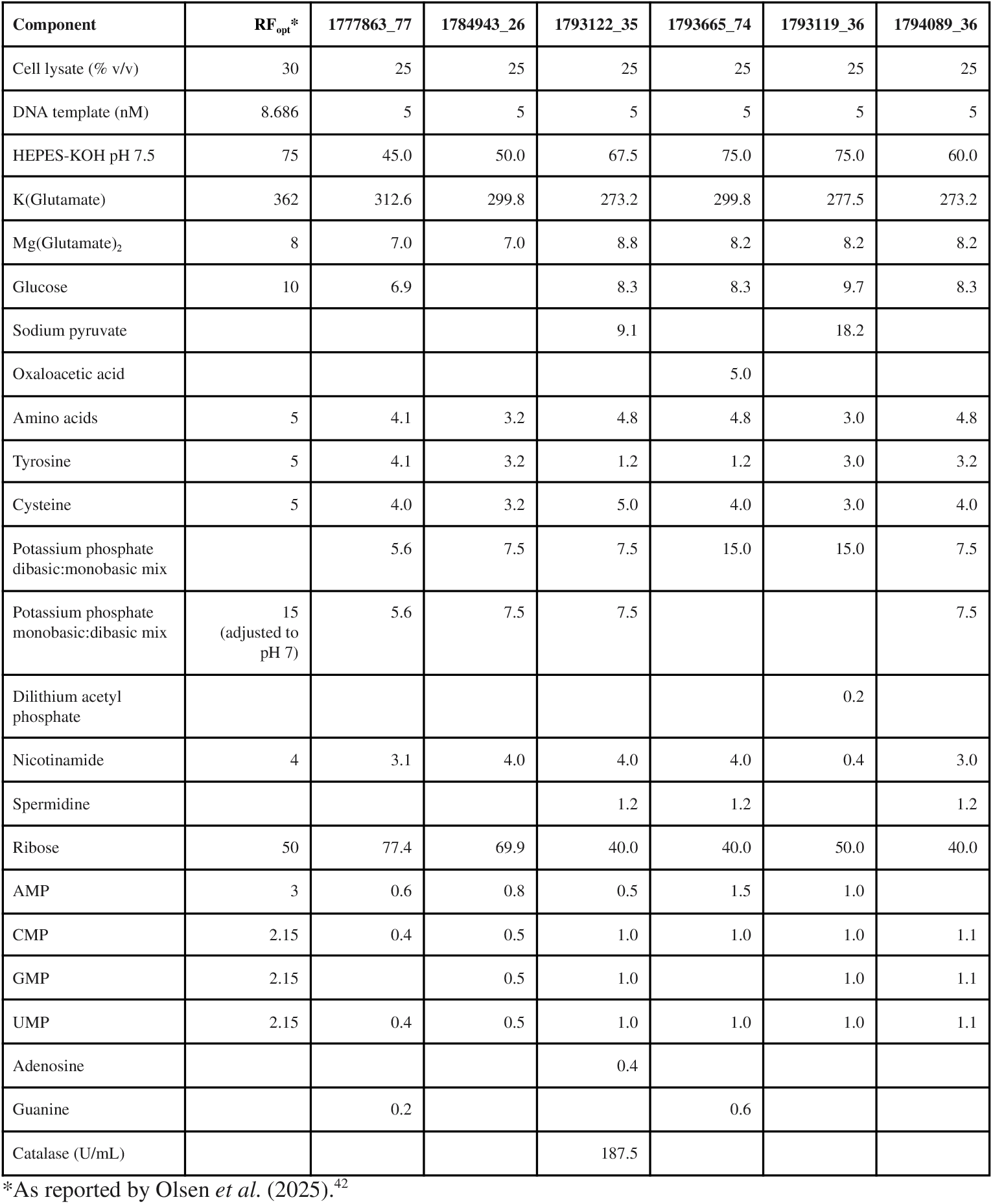
Top performing CFPS reaction compositions identified in this work. All concentrations are in mM, unless otherwise noted. See Supplemental Table 1 for chemical abbreviations and compositions of stock solutions.

| Component | RF<sub>opt</sub>* | 1777863_77 | 1784943_26 | 1793122_35 | 1793665_74 | 1793119_36 | 1794089_36 |
| --- | --- | --- | --- | --- | --- | --- | --- |
| Cell lysate (% v/v) | 30 | 25 | 25 | 25 | 25 | 25 | 25 |
| DNA template (nM) | 8.686 | 5 | 5 | 5 | 5 | 5 | 5 |
| HEPES-KOH pH 7.5 | 75 | 45.0 | 50.0 | 67.5 | 75.0 | 75.0 | 60.0 |
| K(Glutamate) | 362 | 312.6 | 299.8 | 273.2 | 299.8 | 277.5 | 273.2 |
| Mg(Glutamate)<sub>2</sub> | 8 | 7.0 | 7.0 | 8.8 | 8.2 | 8.2 | 8.2 |
| Glucose | 10 | 6.9 |  | 8.3 | 8.3 | 9.7 | 8.3 |
| Sodium pyruvate |  |  |  | 9.1 |  | 18.2 |  |
| Oxaloacetic acid |  |  |  |  | 5.0 |  |  |
| Amino acids | 5 | 4.1 | 3.2 | 4.8 | 4.8 | 3.0 | 4.8 |
| Tyrosine | 5 | 4.1 | 3.2 | 1.2 | 1.2 | 3.0 | 3.2 |
| Cysteine | 5 | 4.0 | 3.2 | 5.0 | 4.0 | 3.0 | 4.0 |
| Potassium phosphate dibasic:monobasic mix |  | 5.6 | 7.5 | 7.5 | 15.0 | 15.0 | 7.5 |
| Potassium phosphate monobasic:dibasic mix | 15<br>(adjusted to pH 7) | 5.6 | 7.5 | 7.5 |  |  | 7.5 |
| Dilithium acetyl phosphate |  |  |  |  |  | 0.2 |  |
| Nicotinamide | 4 | 3.1 | 4.0 | 4.0 | 4.0 | 0.4 | 3.0 |
| Spermidine |  |  |  | 1.2 | 1.2 |  | 1.2 |
| Ribose | 50 | 77.4 | 69.9 | 40.0 | 40.0 | 50.0 | 40.0 |
| AMP | 3 | 0.6 | 0.8 | 0.5 | 1.5 | 1.0 |  |
| CMP | 2.15 | 0.4 | 0.5 | 1.0 | 1.0 | 1.0 | 1.1 |
| GMP | 2.15 |  | 0.5 | 1.0 |  | 1.0 | 1.1 |
| UMP | 2.15 | 0.4 | 0.5 | 1.0 | 1.0 | 1.0 | 1.1 |
| Adenosine |  |  |  | 0.4 |  |  |  |
| Guanine |  | 0.2 |  |  | 0.6 |  |  |
| Catalase (U/mL) |  |  |  | 187.5 |  |  |  |

\*As reported by Olsen *et al.* (2025).<sup>42</sup>

The autonomous lab was able to improve upon SOTA by producing superfolder green fluorescent protein (sfGFP) at a specific cost of $422/g versus Olsen *et al*.’s report of $698/g (Table 1). The composition obtained from the autonomous lab reduces CFPS specific cost by 40% and increases protein titers by 27% relative to SOTA. Although reagents only cost was not an explicit model objective, we also reduced reagent costs from $60/g to $26/g, a reduction of 57% over SOTA. Note that although the preprint published by Olsen *et al*. on August 1, 2025 quoted a reagents only cost of $55/g, the authors subsequently updated this cost to $60/g prior to final publication (Michael Jewett, personal communication).

### We used a Pydantic model to filter GPT-5-designed experiments for scientific validity

A key technical hurdle in the use of autonomous labs for scientific experimentation is ensuring that AI models generate valid experimental designs that can be physically performed on the available equipment. To address this hurdle, we made use of Pydantic, a Python data validation library, to formalize the shared contract for experimental design, encode physical and logistical constraints, and define the schema from which Catalyst protocols are generated and experimental results are returned and analyzed. We imposed key experimental design constraints including plate format and layout, standards, controls, replication factor, and allowed reagent volumes. We also parameterized the available CFPS reagents, stock concentrations, chemical compositions and unit costs (Supplemental Table 1).

GPT-5 interacted with the Pydantic schema to produce a JSON file representation of a “CFPS experiment,” a 384-well plate with specified reaction compositions. The Pydantic schema ensures that GPT-5 produces an experimental design that is both syntactically well-formed for Catalyst protocol generation and physically executable in Ginkgo’s cloud lab. The plate JSON file serves as a convenient abstraction and interface between the LLM, which is focused on experimental intent and design, and the cloud laboratory, which handles all the complexities of physical implementation of the experiment.

### GPT-5 wrote useful lab notebook entries documenting its scientific reasoning

As discussed above, GPT-5’s lab notebook entries summarized prior data, interpreted results and proposed new hypotheses for testing. In particular, GPT-5 suggested new reagents that it deemed potentially useful to reduce specific cost, accompanying rationale for their inclusion and target concentration ranges to try (Supplemental Material section 7.5). For instance, GPT-5 noted that nucleotide triphosphates (NTPs) were a dominant cost in CFPS reactions and prioritized reagents that it reasoned could support NTP regeneration such as polyphosphate, acetyl phosphate, and potassium phosphate or NTP synthesis such as nucleotide monophosphates (NMPs), nucleosides, bases and ribose, in advance of being provided the Olsen *et al*. (2025) preprint. Thus, GPT-5 anticipated certain reagents such as NMPs, potassium phosphate and ribose that were critical to the reagent cost advances described by Olsen *et al*. Note that GPT-5 did not list nicotinamide in its prioritized reagents to add; however, GPT-5 did try nicotinamide and ultimately used it in all top-performing reaction compositions of this work (Table 2). Of the more than 20 additional reagents proposed by GPT-5, several were components of one or more of our top-performing reaction compositions, including NMPs, glucose, potassium phosphate, dilithium acetyl phosphate, adenosine, guanine, catalase, and oxaloacetic acid. The model also suggested various enzymes that it reasoned could be helpful in NTP regeneration or synthesis, but most of these could not be sourced commercially and were not included.

### Reaction designs improved significantly with computer tools, DNA template and cell lysate improvements, experimental (meta)data return and the SOTA report

We provided GPT-5 with access to the internet, a computer, and data analysis packages as well as the preprint on SOTA by Olsen *et al*. (2025)^42^ (including supplementary materials) starting in *Step 3*. Since GPT-5 had access to a computer, we also chose to provide it with access to the Pydantic model itself so that we could return additional experimental (meta)data for the model to consider in its experimental design. Thus, in this setting, GPT-5 was able to combine insights from the SOTA report with previous experimental results and other online publications (see References section of the GPT-5 generated lab notebook entry in Supplementary Materials section 7.3), and design CFPS experiments accordingly. Together, these changes along with cell lysate and DNA template improvements (Supplementary Materials section 7.4) resulted in a performance jump in both specific cost and protein titer (Figure 2). In fact, the model achieved many reactions with better specific cost than SOTA for 384-well plates in *Step 3*. This performance jump is consistent with prior work by OpenAI demonstrating significant model performance gains with tool use across multiple benchmarks.^44^

### Heuristic scores from grading of experimental designs did not obviously correlate with superior designs

At each step, we ranked and selected all Pydantic-validated GPT-5 generated experimental designs using one of three scoring methods: (A) “heuristic” in which another LLM with a “principal investigator” persona critiqued the design and justification (Supplementary Material section 7.6); (B) “random” in which only Pydantic model validation and no scoring was used; and (C) “rubric” in which GPT-5’s reasoning process was graded against a rubric crafted by human experts to encourage thorough analysis and reaction diversity (Supplementary Material section 7.7). The scoring used was: *Steps 0-2*, heuristic; *Steps 3-4*, random; *Step 5*, rubric. We did not conclusively establish whether heuristic scoring correlated with superior designs since (A) top-performing reaction compositions came from designs with both high and low heuristic scores and (B) improvements in specific cost occurred even in the absence of scoring (*Step 3* and *Step 4*).

### GPT-5 discovered novel, improved CFPS reaction compositions

In six steps spanning six months, GPT-5 was able to find CFPS reaction compositions that surpassed state of the art in terms of both specific cost ($/g) and titer (g/L) (Table 1). The top performing reaction compositions differ from SOTA both in reaction composition and component concentration (Table 2).

When prompted to synthesize insights from *Steps 3-5*, GPT-5 (guided by human input and subsequently checked with our own median titer calculations) made several observations and potential rationales that we found plausible but require further experimental validation.

1. Adding cheap HEPES buffer has an outsized impact on specific cost by preventing yield collapse from pH changes. Generally, it found that higher HEPES concentrations increased titer and thus decreased cost. GPT-5 also noted that there is an interdependence between other reaction components and the optimal amount of buffer. For example, in *Step 4*, reactions without added HEPES-KOH pH 7.5 (e.g., just the HEPES in the base buffer) had a median titer of 0.057 g/L (95% confidence interval 0.053–0.061), whereas reactions with an added 1200-2000 nL HEPES-KOH pH 7.5 had a median titer of 0.177 g/L (95% confidence interval 0.160–0.222).
2. Phosphate must be buffered, with mono- and dibasic potassium phosphates within an optimum concentration and pH range, lest titers fall precipitously. For example, in *Step 5*, reactions with either no potassium phosphate or very high potassium phosphate (> 25 mM) had very low titer or approximately zero titer, respectively; whereas concentrations of 5-10 mM produced strong titers. This observation is consistent with our result that all top-performing reaction compositions in this work as well as RF_opt_ from Olsen *et al*. (2025) include potassium phosphate (Table 2).
3. In concurrence with Olsen et al.,^42^ GPT-5 concluded that use of NTPs is not a cost-effective route to high protein titers and that NMPs yield a better specific cost. The model also noted that NMPs cannot be fully replaced by the corresponding nucleosides or bases, but partial replacement does reduce costs while preserving titers. Again, this finding agrees with Olsen et al., who note that GMP may be replaced by guanine and ribose without loss of titer, a result we confirmed (Tables 1 and 2, reaction compositions 1777863_77 and 1793665_74). GPT-5 also found that AMP may be partially replaced by adenosine (Tables 1 and 2, reaction composition 1793122_35). In *Step 5*, reactions with adenosine had a median titer of 0.279 (95% confidence interval 0.273–0.285) and those without had a median titer of 0.138 (95% confidence interval 0.136–0.139). However, attempts to replace CMP and UMP with their pyrimidine nucleoside counterparts failed.
4. Spermidine is a low-cost reaction component for which addition correlates with increased protein titers. Specifically, in *Step 5*, reactions with spermidine had a median titer of 0.253 (95% confidence interval 0.248–0.259) whereas those without had a median titer of 0.131 (95% confidence interval 0.129–0.133). GPT-5 asserted that polyamines such as spermidine stabilize nucleic acids and ribosomal function, thereby improving transcription and translation efficiency.
5. Sodium hexametaphosphate did not have a beneficial effect on titers.
6. Reaction costs are dominated by cell lysate and DNA template. But improvements in titer from buffering / pH control or from addition of spermidine can materially improve the overall specific cost.

To evaluate whether the CFPS reaction compositions developed here could be used to produce other proteins of interest, we tested CFPS reaction composition 1793119_36 with a panel of twelve proteins (each of which had a C-terminal HiBiT complementation tag) and sfGFP. We found that six of twelve proteins were produced at sufficiently high titer to be visible by SDS-PAGE (Supplemental Figure 4) in addition to sfGFP and nine of twelve proteins were detectable by HiBiT assay (data not shown), suggesting that further CFPS optimization may be needed for particular proteins of interest.

## 3. Discussion

Our work demonstrates the real-world application of an autonomous lab to address an open scientific problem, the simultaneous cost- and titer-optimization of cell-free protein synthesis. We used OpenAI’s GPT-5 to plan and design experiments, and Ginkgo’s cloud laboratory, based on RAC automation technology, to execute those experiments. Human intervention was primarily limited to the preparation, loading and unloading of reagents and consumables. We were able to improve upon the prior state of the art to produce sfGFP at a specific cost of $422/g versus $698/g, representing a 40% reduction in reaction cost per gram of protein produced over SOTA. Although the CFPS reaction costs quoted here are not directly comparable to the prices of commercially-available CFPS kits, it is notable that the NEBExpress kit, as one example, has a list price of $800,000/g of protein produced based on reported specifications.^45^

Importantly, our GPT-5-driven autonomous lab was able to reduce reaction costs while *increasing* CFPS titer. Use of reaction composition 1793122_35, for example, improved protein titer by 27% in 2-mL reaction tubes (3.04 vs. 2.39 g/L; Table 1), and by 188% — nearly 3-fold higher — in 384-well plate reactions (0.72 vs. 0.25 g/L; Table 1). We anticipate that additional optimization for (A) particular reaction geometries, (B) particular reaction volumes, or (C) the protein of interest could further improve *E. coli* CFPS protein production.

Use of OpenAI’s GPT-5 for experimental planning and design and Ginkgo’s RAC-based automated cloud laboratory for experimental execution enabled data generation at a scale sufficient to support reinforcement learning. In total, we ran over 580 microtiter plates, tested 36,000 reaction compositions, and generated almost 150,000 datapoints in six months, with *Steps 3–5* taking just two months. We believe this scale and speed of successful experimentation would not have been possible without the use of an LLM-driven autonomous lab.

In autonomous labs, experimental costs are dominated by reagents and consumables, not personnel. We believe that our use of an autonomous lab to optimize CFPS is a harbinger of things to come. Particularly apropos, one of the many potential applications of this work is a cloud laboratory running low-cost CFPS workflows as a useful complement to the many AI-based protein design tools available today.^46–51^

We chose not to use GPT-5 to write the Catalyst protocols responsible for executing the RAC-based workflows. Indeed, all Catalyst protocols were written by Ginkgo personnel. Instead, we constrained GPT-5, via the Pydantic schema, to produce experimental designs within a predefined operational envelope of Ginkgo’s cloud laboratory. We believe that this constraint appropriately focused the GPT-5-driven autonomous lab on exploring relevant regions of biological parameter space rather than on writing, attempting to run, and debugging RAC-compatible protocols. We intend our work to open the aperture of what Thomas Kuhn calls “normal science”^52^ that can be successfully performed using an LLM-driven autonomous lab.

A common concern with the use of the current generation of LLMs for scientific research is the possibility of hallucinations. In part due to our use of the Pydantic schema to validate experimental designs produced by GPT-5, we found that hallucinations were quite rare: less than 1% of the executed designs were flawed (just two of 480 total 384-well plates designed by GPT-5 across *Steps 0–5*). In one plate design, the model attempted to overwrite the required volume of 2 µL for 10X base CFPS buffer (see Materials and Methods) so that it could add additional volumes of other available reagents while remaining within the 20-µL total volume specification, resulting in a total CFPS reaction volume above our 20-µL specification. We updated the validation in the Pydantic schema to prevent this error from recurring. In a later design, GPT-5 wrote code that had a minor bug (converting nL to μL by dividing by 10^6^), resulting in a plate where each well contained only glucose and ribose. Unsurprisingly, in the absence of nucleic acid building blocks, these CFPS reactions produced no protein. Overall, given that the model designed over 480 plates and we only discovered 2 plates with fundamental design flaws post execution, the autonomous lab designed and executed scientific experiments remarkably well.

In the immediate future, we believe work such as that described here—optimizing CFPS reaction cost and titer using LLM-driven autonomous lab experimentation—will become common across a wide range of biological, biomedical and even chemical science. And just a few years out, as reasoning models grow ever more powerful, and modular laboratory automation becomes pervasive, autonomous lab experimentation will increasingly supplant traditional labwork, allowing scientists to focus on harder and more fulfilling work than designing and executing experiments.

## 4. Materials and Methods

### Pydantic model

The Pydantic (v2.11.7; https://docs.pydantic.dev/) schema comprised three primary models to define the interface between experimental design by GPT-5,^41^ a reasoning model released by OpenAI on August 7, 2025, and execution on Ginkgo’s cloud lab. The Plate model represented the complete experimental design, including plate geometry, plate layout, sample definitions, and high-level composition metadata such as reserved columns and the number and placement of controls. The Sample model represented an individual cell-free protein synthesis reaction, capturing the experimental condition at the well level, validating the desired replication for each sample. The ReagentList model specified the set of reagents and their associated volumes for each sample; at this level, the availability of reagents and permissible reagent volumes were validated against inventory and constraint rules. Together, these models provided a structured, machine-interpretable specification of experimental designs that could be reliably transformed into executable Catalyst protocols. For each iterative step, we generated 2X the target number of designs from GPT-5 because not all experiment definitions passed Pydantic validation. The Pydantic model used in this work has been open-sourced (Supplementary Material section 7.8).

### Cell lysate preparation

Lysate was generated from a fermented *E. coli* (BL21 Star (DE3)) strain. We grew cells overnight at 30°C in LB media to prepare starter culture. The following day, we combined all starter cultures and confirmed OD. We aseptically charged inoculum to Ginkgo’s FM39 production medium supplemented with isopropyl-β-D-thiogalactopyranoside (IPTG) to induce T7 RNA polymerase expression in 5L Bioflo 320 vessels (Eppendorf) for fed-batch style fermentation with temperature, oxygen, pH, glucose and antifoam control. Once fermentation was complete, we harvested and pelleted cultures. We discarded the supernatant and washed the pellet three times in the harvest/lysis buffer. After the third wash the pelleted material was flash frozen in liquid nitrogen and stored at -80°C.

We removed the washed fermentation pellet from the freezer and thawed at room temperature. We supplemented the harvest/lysis buffer with DTT to the thawing pellet and agitated it to ensure a homogenous mixture. Once fully thawed and homogenous, we lysed the material with a PandaPLUS 2000 homogenizer (GEA). We clarified the resulting lysate twice via centrifugation. We then aliquoted, flash froze and stored the final clarified lysate at -80°C.

### Plasmid DNA preparation

Plasmid m9134733 (length of 2511 base pairs; molar mass of 1,546,533 g/mol) was synthesized and cloned at Ginkgo using Type IIS restriction enzyme-based DNA assembly and standard molecular cloning into *E. coli* DH5alpha strain (Supplementary Material section 7.9). The plasmid sequence was based on pJL1-sfGFP (Addgene plasmid #69496; http://n2t.net/addgene:69496)^30^ used in state of the art by Olsen *et al*. (2025).^42^ We introduced minor sequence modifications as a byproduct of our DNA assembly process at the DNA assembly junctions. The plasmid insert, harboring the sfGFP with C-terminal Strep-tag II transcription unit including T7 promoter, ribosome binding site, coding sequence and terminator, remained unchanged from pJL1-sfGFP. We outsourced giga-scale plasmid DNA production (from single colonies on LB agar plates with 50 µg/mL kanamycin) to Genewiz (Azenta Life Sciences).

### Cell-free protein synthesis reactions

Cell-free protein synthesis reaction compositions were high-throughput end-to-end screened using fluorescence plate reader measurement-compatible 384-well plates (Greiner, cat. #781209; surface area to volume ratio: 1 mm_2_/mm_3_). Low-throughput, follow-up assessments of cell-free protein synthesis reaction oxygen dependency during incubation were performed using intermediate 96-well plates (Thermo Fisher Scientific, cat. #95040450; surface area to volume ratio: 3 mm_2_/mm_3_) and the 2-mL tubes (Axygen, cat. #14-222-180; surface area to volume ratio: 5 mm_2_/mm_3_) used in Olsen *et al*. (2025).^42^ Cell-free protein synthesis reactions that completed incubation were then transferred to the 384-well format plates, for downstream fluorescence plate reader measurements.

Unless otherwise noted, all cell-free protein synthesis reactions were performed in a total volume of 20 µL, and contained 25% v/v cell lysate, 5 nM plasmid DNA template and 2 µL of “10X base CFPS buffer” (300 mM HEPES, pH 7.5; 1.5 M potassium glutamate, 26 mM magnesium glutamate, 10 mM 17 amino acid mix [all 20 canonical amino acids except glutamate, tyrosine and cysteine], 10 mM tyrosine, pH 12; 10 mM cysteine), used to establish a buffer, salts and amino acid baseline in all reactions using more concentrated stock solutions, thus ensuring sufficient leftover volume (11 µL) for acoustic dispensing of the remaining variable reaction components. All essential and non-essential variable reaction components (additional buffers and salts, nucleobases, nucleosides, nucleotides, amino acids, ATP regeneration components, co-factors, solvents, detergents etc.) are detailed in Supplemental Table 1.

Cell-free protein synthesis reactions were set up, incubated and measured using Ginkgo’s cloud laboratory built from RAC lab automation technology (Supplemental Figure 2) and Catalyst software (Supplementary Figure 1 and 3). Variable reaction components were acoustically dispensed, using the Echo 525 acoustic dispenser (Beckman Coulter), from acoustic dispensing-compatible 384-well source plates (Beckman Coulter, cat. #001-14555) to columns 4-24 of the fluorescence plate reader measurement-compatible 384-well destination plates (Greiner, cat. #781209). Based on offline, manual liquid handling experiments, different liquid classes were used for different reaction components: the 384PP_AQ_BP liquid class was used for all reagents except for 60% v/v DMSO and 10% v/v Brij-35, for which the 384PP_AQ_SPHigh liquid class was used. Each 384-well source plate was unique and dedicated to one 384-well destination plate. Each 384-well destination plate was pre-dispensed with 2 µL 10X base CFPS buffer into all columns, using the MultiFlo FX bulk dispenser (Agilent). The first three columns of each 384-well destination plate were dedicated to sfGFP protein standards and did not receive any reagents; instead, they were backfilled with 11 µL 1X Phosphate-Buffered Saline, pH 7.5 (Teknova, cat. #P5275), using the MultiFlo FX bulk dispenser (Agilent).

Next, the sfGFP protein standards and plasmid DNA template were liquid transferred (constant 2 µL volume stamp), using the Bravo 384 liquid handler (Agilent), from a pre-made, reusable 384-well source plate (Greiner, cat. #781281) into the pre-filled (with variable reaction components) 384-well destination plate. The reaction setup was completed by a cell lysate bulk dispense (5 µL into all columns), using the MultiFlo FX bulk dispenser (Agilent), followed by a brief (30 seconds) linear shake on the instrument.

Quality control fluorescence plate reader measurements were taken at ambient temperatures (top read, averaging 10 flashes per well using standard settings with 485 nm excitation and 520 nm emission), before the start of cell-free protein synthesis reaction incubations, using the PHERAstar FSX plate reader (BMG Labtech), to confirm the presence of sfGFP protein standards and successful addition of cell lysate. The reactions were generally incubated at 30°C for 20 hours without shaking using the SteriStore (HighRes Biosolutions) though during periods of high Ginkgo cloud lab utilization, incubation times were sometimes extended up to 30 hours. Depending on instrument availability, some plate sets were instead incubated at 30°C for 6 hours, with 400 rpm shaking, using the Cytomat 2 C-LiN (Thermo Fisher Scientific). Non-breathable, aluminum seals were used to seal the 384-well destination plates. Follow up assessments in 96-well plates were incubated at 30°C at 400 rpm with a 3mm orbital throw for 20 hours, using the Cytomat 2 C-LiN (Thermo Fisher Scientific). Follow up assessments in 2-mL tubes were incubated at 30°C at 400 rpm with a 3mm orbital throw for 20 hours, using a Multitron shaking incubator (Infors HT).

### Protein Titer Quantification

Finished cell-free protein synthesis reactions were also processed using Ginkgo’s cloud laboratory. 60 µL of 1X Phosphate-Buffered Saline, pH 7.5 (Teknova, cat. #P5275) was bulk dispensed into each well of the 384-well destination plate, using the MultiFlo FX bulk dispenser (Agilent) to reduce well-to-well variation in measured fluorescence. Lower-throughput, follow-up assessments of cell-free protein synthesis in intermediate 96-well plates and 2-mL tubes required dynamic adjustment of dilution factor, based on the expected protein titer range and our standard curve range, i.e. variable volumes of 1X Phosphate-Buffered Saline, pH 7.5 were added to the intermediate plate wells / tubes and 80 µL of each diluted reaction were transferred into the 384-well destination plates for fluorescence measurement. The varying dilution factors were accounted for in the final protein titer calculations. Final fluorescence measurements were taken with the Spark plate reader (Tecan) at ambient temperatures (top read, 485 nm excitation with 20 nm bandwidth, 535 nm emission with 20 nm bandwidth, measurement averages 30 flashes per well). The output relative fluorescence unit data was immediately available for downstream computer analysis, upon a finished plate read Catalyst software “event”. Protein titers were calculated using standard curve interpolation with RANSAC exclusion of outliers, using either linear or log transformed values to minimize the mean absolute percentage error (MAPE) of the fit curve. The standard curve was set up using 15 different concentrations in technical triplicate. Their reaction concentrations are detailed in Supplemental Table 2.

We made the sfGFP protein standards in-house, using the m9134733 plasmid DNA template. The sfGFP protein expression was driven by a T7 promoter expression system in *E. coli* (BL21 (DE3) strain). We induced protein expression at mid-log phase with 0.2 mM IPTG in a shake flask liquid culture, grown in TB containing 0.4% v/v glycerol and 50 µg/mL kanamycin at 37°C. We harvested cells via centrifugation, after 24 hrs of growth in the liquid cell culture, mechanically lysed using the PandaPLUS 2000 homogenizer (GEA), and filtered with a 0.2 µm filter. The sfGFP protein, harboring a C-terminal Strep-tag, was purified via affinity chromatography. We loaded 50 mL of the clarified cell lysate onto a 5 mL Strep-Tactin XT FF column (Cytiva) on an ÄKTA Avant system (Cytiva), equilibrated with a running buffer (50 mM HEPES, 150 mM NaCl, pH 8.0). We eluted the sfGFP protein with an elution buffer (50 mM HEPES, 150 mM NaCl, 50 mM biotin, pH 8.0). We concentrated eluate and buffer-exchanged into a 20 mM HEPES, pH 7.4 buffer, using tangential flow filtration with a 10 kDa mPES TFF cassette (Repligen). We determined protein concentration via an absorbance measurement at 280 nm, using the calculated extinction coefficient. Protein aliquots were stored at −80°C.

## 5. Acknowledgments

## Author Contributions

OpenAI: Jerry Tworek, Ahmed El-Kishky and Tejal Patwardhan conceived of the study. Ahmed El-Kishky, Mostafa Rohaninejad, and Tejal Patwardhan provided technical guidance. Yunxin Joy Jiao and Edmund L. Wong ran the GPT-5 plate design experiments and related analysis.

Ginkgo Bioworks: Reshma P. Shetty, Paulina Kanigowska, Jose E. Cortez and Michal Jastrzebski conceived of the study. Elizabeth A. Gendreau and Alexus A. Smith wrote the Catalyst protocols for cell-free protein synthesis workflows. Brad A. Chapman, Jose E. Cortez, Ryan Harry, and Michal Jastrzebski wrote the Pydantic model and the software code to connect model-generated Pydantic code to the Ginkgo cloud laboratory. Bradley P. Barber, Trinh Thanongsinh, Andrew Nesson, Bibek Lama, Brandon Nichols, Cameron LaFrance, and Tenzing Nyima synthesized and cloned the plasmid DNA template. José C. Morales Hemuda, Ian Graves, Rahul Karandikar, Christopher Lionetti, Kevin Christopher, Andrew L. Consiglio, Isis Botelho Nunes da Silva, and Alvaro R. Bautista-Ayala prepared various cell lysate batches. Kevin Christopher and Robert H. Dahl prepared purified sfGFP protein standards. Robert H. Dahl, James Dooner, William McCusker, Duy X. Nguyen and Alyssa Tran prepared reagent stock solutions. Alexus A. Smith, Ronan C. Donovan, Pooyan Tirandazi, Robert H. Dahl, Paulina Kanigowska, Reshma P. Shetty, Austin J. Che and Nathan C. Tedford prepared, loaded and unloaded reagents and consumables onto and off of the RAC system during experimental execution and performed all manual experiments noted. Ryan Harry, Brad A. Chapman, Christopher J. Bremner, Michael McDuffie, and Sean Atkins provided software assistance for experimental execution on the Ginkgo cloud laboratory. Elizabeth A. Gendreau, Alicia Byrn, Alycia Wong, Lizvette Ayala-Valdez, Bryan Cai and Rashard Thornhill provided remote monitoring support for the Ginkgo cloud laboratory. Will Serber conceived of RAC automation technology underpinning the Ginkgo cloud laboratory. Daniel Smith, Thomas F. Knight, Jr. and Walter Thavarajah provided scientific guidance on cell-free protein synthesis. Monica P. McNerney provided technical guidance on cell lysate preparation.

Reshma P. Shetty, Yunxin Joy Jiao, David Borhani, Paulina Kanigowska, Edmund L. Wong and Christopher Lionetti wrote the paper.

We thank Sriram Kosuri and Joseph H. Davis for valuable comments on the manuscript.

