## Supplementary Materials for "Using a GPT-5-driven autonomous lab to optimize the cost and titer of cell-free protein synthesis"

### 7.1 Supplemental Tables

*Supplemental Table 1: Reagents used in this work.* Reagent costs are based on those reported by Olsen *et al.* (2025), where possible, for comparison of reaction costs obtained in this work to prior state of the art (SOTA).[42]

| Reagent | Vendor & Catalog # | Stock Concentration | Units | Components | Cost per mL |
|---|---|---|---|---|---|
| **Core Components** | | | | | |
| Cell lysate | N/A | 1 | X | *E. coli* (BL21 STAR (DE3)) strain | $4.000 |
| DNA template | N/A | 0.050 | M | Plasmid m9134733 encoding sfGFP | $1.933 |
| **Diluent** | | | | | |
| Nuclease-free water** | Invitrogen 10-977-023 | N/A | N/A | Nuclease-free water | $0.000 |
| **Core Salt & Buffer Components** | | | | | |
| Potassium glutamate K(Glu) | Sigma G1501 | 0.875 | M | L-Glutamic acid potassium salt monohydrate | $0.070 |
| Magnesium glutamate $Mg(Glu)_2$ | Sigma 49605 | 0.500 | M | L-Glutamic acid hemimagnesium salt tetrahydrate | $0.065 |
| HEPES-KOH pH 7.5 | Boston Bioproducts BBH-75-K | 1 | M | HEPES, potassium hydroxide | $0.130 |
| **Core Transcription Components** | | | | | |
| Adenosine 5′-triphosphate (ATP) | Sigma A2383 | 0.100 | M | Adenosine 5′-triphosphate disodium salt hydrate, potassium hydroxide | $1.539 |
| Cytidine 5′-triphosphate (CTP) | Sigma C1506 | 0.100 | M | Cytidine 5′-triphosphate disodium salt, potassium hydroxide | $27.937 |
| Guanosine 5′-triphosphate (GTP) | Sigma G8877 | 0.100 | M | Guanosine 5′-triphosphate sodium salt hydrate, potassium hydroxide | $39.343 |
| Uridine 5′-triphosphate (UTP) | Sigma U6625 | 0.100 | M | Uridine 5′-triphosphate trisodium salt hydrate, potassium hydroxide | $37.846 |
| **Core Translation Components** | | | | | |
| 17 amino acid mix | See below | 0.050 | M | see below* | $0.083 |
| Tyrosine pH 12 | Sigma T8566 | 0.050 | M | L-Tyrosine, potassium hydroxide | $0.011 |
| Cysteine | Sigma C7352 | 0.200 | M | L-Cysteine | $0.024 |

| Reagent | Vendor & Catalog # | Stock Concentration | Units | Components | Cost per mL |
|---|---|---|---|---|---|
| **Core ATP Regeneration Components** | | | | | |
| Sodium pyruvate | Sigma P2256 | 100 | g/L | Sodium pyruvate | $0.164 |
| Maltose | Sigma M5895 | 50 | g/L | D-(+)-Maltose monohydrate | $0.020 |
| Ribose | Sigma R7500 | 100 | g/L | D-(-)-Ribose | $0.180 |
| Maltodextrin 17 | Sigma 419699 | 300 | g/L | Maltodextrin , dextrose equivalent 16.5 - 19.5 | $0.122 |
| **Co-factors** | | | | | |
| Folinic acid | Sigma TA9H98774066 | 10 | g/L | Folinic acid, potassium hydroxide | $8.978 |
| NAD | Sigma N6522 | 0.100 | M | β-Nicotinamide adenine dinucleotide, potassium hydroxide | $33.348 |
| cAMP | Sigma A9501 | 0.200 | M | Adenosine 3′,5′-cyclic monophosphate, potassium hydroxide | $7.480 |
| Spermidine | Sigma S0266 | 0.250 | M | Spermidine | $1.045 |
| **Detergents** | | | | | |
| Brij-35 | Ricca Chemical 1145-32 | 10 | % v/v | Brij® 35 | $0.027 |
| **Solvents** | | | | | |
| Dimethyl sulfoxide (DMSO) | Sigma 472301 | 60 | % v/v | Dimethyl sulfoxide | $0.203 |
| **Additional reagents requested by the model (added after step 1)** | | | | | |
| Pyruvate oxidase (1.5U/mg) | Ag Scientific P-1600-5KU | 10 | g/L | Pyruvate oxidase, Potassium phosphate monobasic, Potassium phosphate dibasic | $48.150 |
| Catalase (>50000U/ml) | Sigma 02071 | 50000 | U/ml | Catalase, Potassium phosphate monobasic, Potassium phosphate dibasic | $5.000 |
| Sodium hexametaphosphate | Sigma 305553 | 0.150 | M | Sodium hexametaphosphate | $0.004 |
| Dilithium Acetyl Phosphate | Sigma A1349332 | 0.050 | M | Dilithium acetyl phosphate, potassium hydroxide | $0.273 |
| Adenosine 5’-monophosphate (AMP) | Sigma 01930 | 0.100 | M | Adenosine 5’-monophosphate disodium salt, acetic acid | $0.699 |
| Cytidine 5’-monophosphate (CMP) | Sigma C1006 | 0.100 | M | Cytidine 5’-monophosphate disodium salt, acetic acid | $1.270 |
| Guanosine 5’-monophosphate (GMP) | Sigma G8377 | 0.100 | M | Guanosine 5’-monophosphate disodium salt, acetic acid | $0.533 |
| Uridine 5’-monophosphate (UMP) | Sigma U6375 | 0.100 | M | Uridine 5’-monophosphate disodium salt, acetic acid | $0.839 |
| Adenosine | Sigma | 0.025 | M | Adenosine, Potassium hydroxide | $0.002 |

| Reagent | Vendor & Catalog # | Stock Concentration | Units | Components | Cost per mL |
|---|---|---|---|---|---|
| | A9251 | | | | |
| Guanosine | Sigma G6264 | 0.025 | M | Guanosine, Potassium hydroxide | $0.012 |
| Cytidine | Sigma C4654 | 0.025 | M | Cytidine | $0.047 |
| Uridine | Sigma U3750 | 0.025 | M | Uridine, Potassium hydroxide | $0.003 |
| Thymidine | Sigma T9250 | 0.025 | M | Thymidine, Potassium hydroxide | $0.108 |
| Adenine | Sigma A8626 | 0.025 | M | Adenine, Potassium hydroxide | $0.010 |
| Guanine | Sigma G11950 | 0.025 | M | Guanine, Potassium hydroxide | $0.003 |
| Cytosine | Sigma C3506 | 0.025 | M | Cytosine, Acetic acid | $0.034 |
| Uracil | Sigma U1128 | 0.025 | M | Uracil, Potassium hydroxide | $0.002 |
| Thymine | Sigma T0376 | 0.025 | M | Thymine, Potassium hydroxide | $0.007 |
| **Additional reagents requested by the model (added after step 2)** | | | | | |
| Glucose | Teknova G0520 | 200 | g/L | Glucose | $0.00222 *Sigma G8270 price to match Olsen *et al.* (2025) |
| Potassium phosphate dibasic | Sigma P3786 | 0.500 | M | Potassium phosphate dibasic | $0.035 |
| Potassium phosphate monobasic | Sigma P0662 | 0.500 | M | Potassium phosphate monobasic | $0.010 |
| Potassium phosphate 1.6:1 dibasic:monobasic (molar ratio) | Sigma P3786 & P0662 | 0.5 | M | Potassium phosphate dibasic, Potassium phosphate monobasic, Potassium hydroxide | $0.024 |
| Potassium phosphate 1.6 :1 monobasic:dibasic (molar ratio) | Sigma P3786 & P0662 | 0.5 | M | Potassium phosphate dibasic, Potassium phosphate monobasic, Potassium hydroxide | $0.018 |
| **Additional reagents (added after step 2)** | | | | | |
| Phosphoenolpyruvic acid cyclohexylammonium salt | Fisher P07581G | 0.1 | M | Phosphoenolpyruvic acid, cyclohexylammonium salt, potassium hydroxide | $2.202 |
| Nicotinamide | Sigma N7559 | 0.1 | M | nicotinamide | $0.002 |
| Base buffer | Individual components listed in table | 10 | X | HEPES, Potassium hydroxide, Arginine, Isoleucine, Aspartic Acid, Alanine, Asparagine, Histidine, Proline, Serine, Threonine, Valine, Leucine, Phenylalanine, Tryptophan, Methionine, Lysine, Glycine, Glutamine, Glacial acetic acid, tyrosine, cysteine, L-Glutamic acid potassium salt monohydrate, L-Glutamic acid hemimagnesium salt tetrahydrate | $0.220 |
| **Additional reagents requested by the model (added after step 4)** | | | | | |

| Reagent | Vendor & Catalog # | Stock Concentration | Units | Components | Cost per mL |
|---|---|---|---|---|---|
| Oxaloacetic acid | Sigma O7753 | 0.5 | M | Oxaloacetic acid, potassium hydroxide, acetic acid | $0.925 |
| Succinic acid | TCI S0100 | 0.5 | M | Succinic acid, potassium hydroxide | $0.004 |
| Potassium formate | Alfa Aesar A14551 | 0.5 | M | Potassium formate, acetic acid, potassium hydroxide | $0.004 |

*17 amino acid mix is composed of Arginine (A8094), Isoleucine (I2752), Aspartic Acid (A7219), Alanine (A7627), Asparagine (A4159), Histidine (H8000), Proline (P0380), Serine (S4500), Threonine (T8625), Valine (V0513), Leucine (L8000), Phenylalanine (P5482), Tryptophan (T0254), Methionine (M9625), Lysine (Santa Cruz sc-207804), Glycine (G7126), Glutamine (G3126) and glacial acetic acid (A6283). All amino acids sourced from Sigma, except where otherwise noted. Catalog numbers in parentheses.

**Nuclease-free water volume needs to be matched with the total volume of other CFPS reagents, to maintain consistent final CFPS reaction volume (20 µL).

*Supplemental Table 2: Concentrations of sfGFP protein standards within the reaction were used for conversion of fluorescence measurements to absolute protein titers.* All concentrations are in µM. The molecular weight of sfGFP protein is 27,015 Da.

| Name | Concentration |
|---|---|
| Standard 1 | 118.90 |
| Standard 2 | 90.00 |
| Standard 3 | 59.45 |
| Standard 4 | 45.00 |
| Standard 5 | 29.73 |
| Standard 6 | 22.50 |
| Standard 7 | 14.86 |
| Standard 8 | 11.25 |
| Standard 9 | 7.43 |
| Standard 10 | 5.63 |
| Standard 11 | 3.72 |
| Standard 12 | 2.81 |
| Standard 13 | 1.86 |
| Standard 14 | 1.41 |
| Standard 15 | 0.93 |

| | |
|---|---|
| Negative control | 0 |

#### 7.2 Supplemental Figures

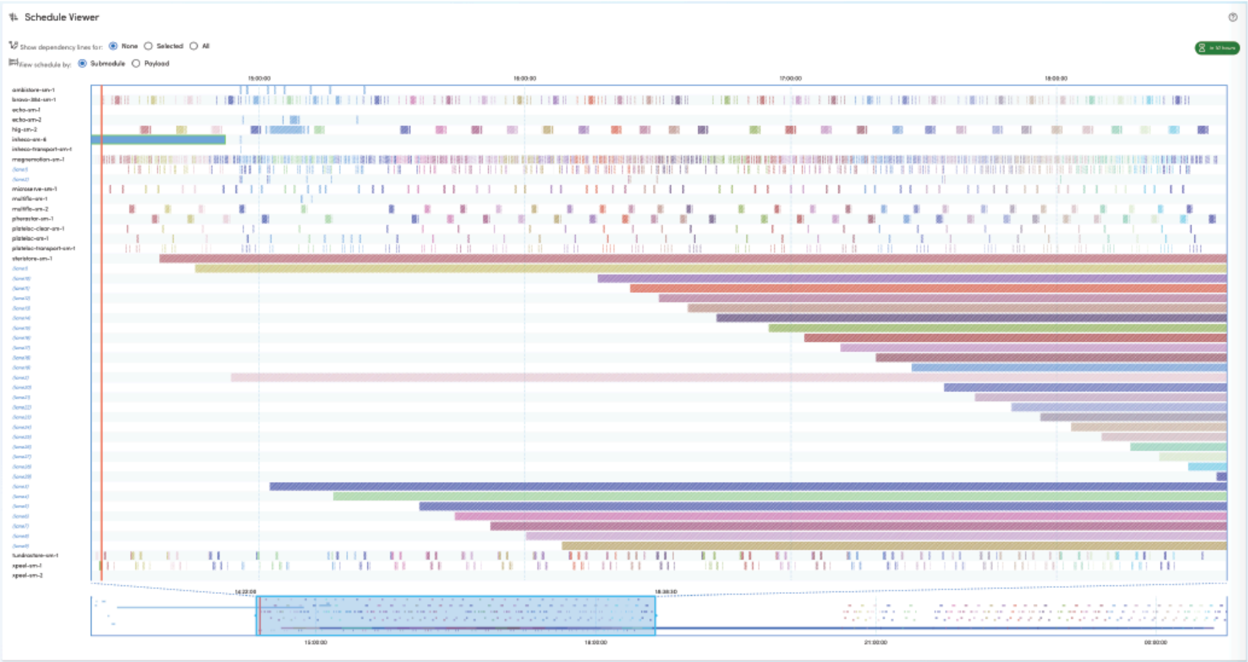


*Supplemental Figure 1: Example schedule of multiple concurrent Catalyst protocols runs for CFPS reaction optimization workflow.* Rows correspond to Ginkgo's cloud lab instruments, within their respective RACs (Catalyst submodules). Each color corresponds to a Catalyst protocol run for a single 384-well destination plate. Each colored rectangle denotes occupancy of a given instrument RAC by a given protocol run action. Some instrument RACs, like the large-capacity static incubator RACs, as well as the sample transport tracks connecting RACs, can support multiple, simultaneous protocol run actions.

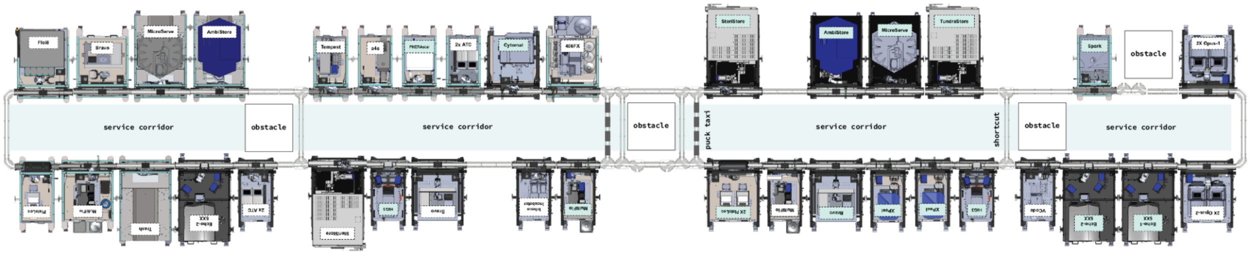


*Supplemental Figure 2: Aerial schematic diagram of Ginkgo's RAC-based cloud laboratory.* Ginkgo's cloud lab comprised thirty-six RACs as of December 31, 2025. CFPS workflows made use of eleven RACs.

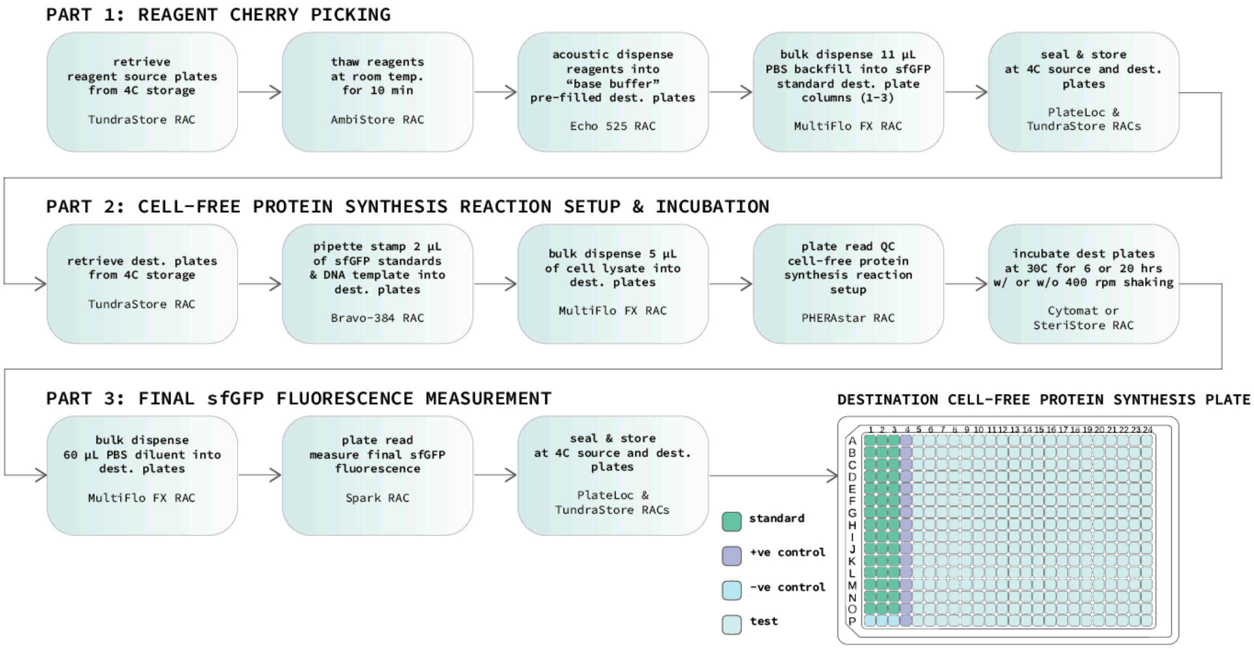


*Supplemental Figure 3: Schematic overview of CFPS reaction optimization workflows as implemented in Catalyst.* We implemented the CFPS workflow as three separate Catalyst protocols for flexibility in workflow execution.

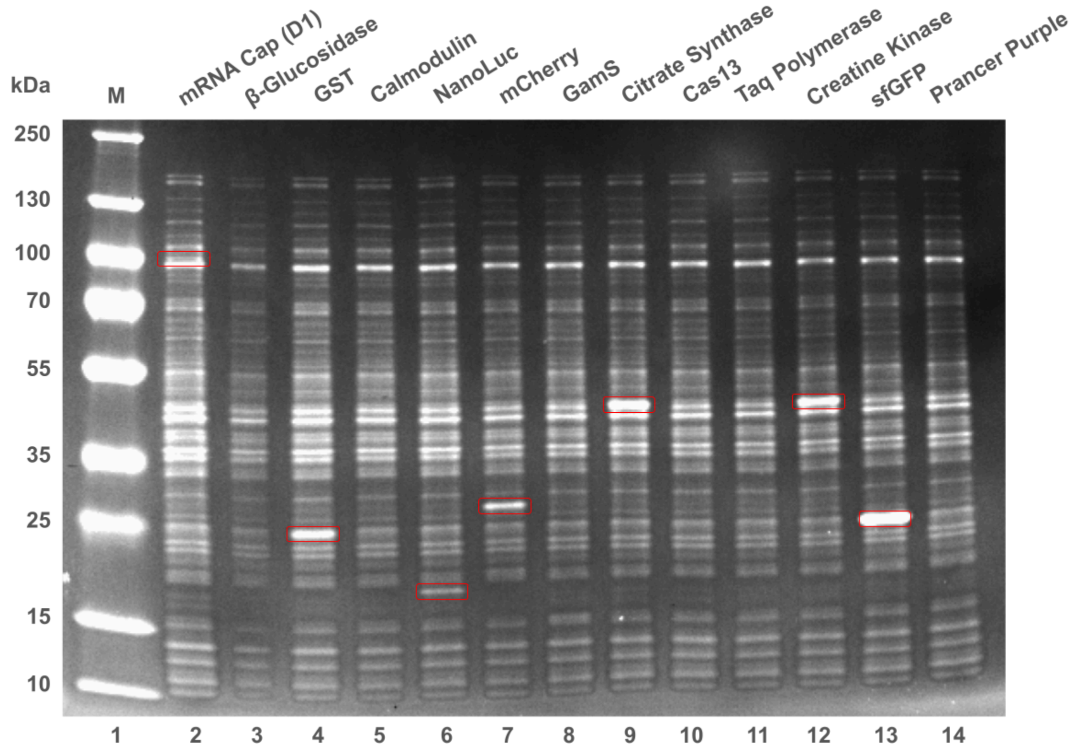


*Supplemental Figure 4: SDS-PAGE gel of twelve proteins plus sfGFP expressed in E. coli CFPS reaction composition 1793119_36. Six of the twelve proteins can be detected by SDS-PAGE.* Twelve proteins (mRNA-capping enzyme D1 subunit, 98.3 kDa, plasmid m8975504; β-glucosidase, 55.3 kDa, plasmid m8975496; glutathione-S-transferase, 28.5 kDa, plasmid m8975528; calmodulin, 18.4 kDa, plasmid m8975485; nanoluciferase, 20.7 kDa, plasmid m8975529; mCherry, 28.3 kDa, plasmid m8975487; GamS, 17.9 kDa, plasmid m8975511; citrate synthase (gltA), 49.6 kDa, plasmid m8975514; Cas13, 133.8 kDa, plasmid m8975476; Taq DNA polymerase, 95.5 kDa, plasmid m8975501; creatine kinase, 44.7 kDa, plasmid m8975477; sfGFP, 27.0 kDa, plasmid m9134733; Prancer purple, 27.1 kDa, plasmid m8975510) were expressed in *E. coli* CFPS reaction composition 1793119_36 at 20 µL volume in 2-mL tubes. CFPS reactions were diluted 10X in 1X Phosphate-Buffered Saline, pH 7.5 (Teknova, cat. #P5275) and frozen. Subsequently, samples were diluted 2X in DNase-free water and then 2X again in Laemmli buffer for a total dilution factor of 40X. Samples were denatured at 95°C for 10 minutes and 3 µL was loaded per lane. For molecular weight assessment, 3 µL PageRuler Plus Prestained Protein Ladder (Thermo Scientific), 10 to 250 kDa was loaded in lane 1 denoted by "M". The gel was run at 180 V for 50 mins, stained with InstantBlue Coomassie (Abcam), washed three times with deionized water, and then imaged on a FluorChem E system (Bio-Techne) using the Coomassie Blue setting. Images were displayed with inverted grayscale and contrast setting of -4.

### 7.3 Example GPT-5-generated lab notebook entry

#### CFPS optimization design – Notes and rationale

This document explains how the 78 CFPS conditions in plate.json were designed to maximize **GFP yield per dollar**. The design integrates (i) the Olsen *et al.* 2025 SOTA study, (ii) analysis of all previous experimental results in data, and (iii) additional literature on reaction energetics and oxygenation. The output plate contains 78 unique conditions (replicates and controls are added downstream by the grader).

---

##### 1. How previous results were analyzed

All raw results in *.json were parsed programmatically. For each run a cfps_models.Plate object was reconstructed, which maps the samples list to well coordinates in plate order. The following data were extracted for each experimental condition:

- **Reagent volumes (nL)** for every reagent used
- **Yield** quantitated as concentration_results['concentration_g_L'][well] (g/L)
- **Cost per reaction** using the reagent field metadata (stock costs are encoded in the cfps_models.ReagentList fields). The Plate.sample_cost property was used, which multiplies ($/mL) × (nL / 1,000,000) and **includes lysate and DNA**. In other words, the cost metric here is *stricter* than Olsen *et al.* (who report the cost of the **reagent mixture** only, not lysate or DNA).

For each unique reagent set (a grouping by the exact reagent list, as pydantic model_dump() values) we calculated:

- **Average yield (g/L)** across replicate wells (plates were randomized; replicate lanes were identified by identical reagent compositions)
- **Cost per liter** = sample_cost × 50,000 (20 µL reactions → 50,000 per liter)
- **g per $** = yield_g_per_L / (cost_per_L) and the reciprocal $ per g

Two classes of experiments were especially informative:

*1.1. 5000 nL lysate, 384-well plates (most comparable to the requested format)*

These runs used 5 µL lysate and 6 h incubations. The best cost-normalized group was obtained in the **RFopt-like** chemistry using low nucleotide monophosphates and polyphosphate:

*Volumes of the best historical group (average 0.219 g/L; $/g ≈ 2596 including lysate/DNA):*

```None
375 nL K-glutamate, 225 nL Mg-glutamate, 300 nL HEPES, 600 nL+600 nL phosphate
6000 nL sodium hexametaphosphate (22.5 mM polyP)
```

```
1500 nL ribose (≈50 mM), 200 nL glucose (≈10 mM), 800 nL nicotinamide (4 mM)
AMP/CMP/UMP 200 nL each (1 mM each) + 800 nL guanine (1 mM) to replace GMP
Amino acids: 1200 nL of 17AA, 1200 nL tyrosine, 200 nL cysteine
```

This condition produced **0.219 g/L** sfGFP at a per-reaction cost of **$0.0114**, so **$/L ≈ $568** and **g per $ ≈ $3.85\times10^{-4}$** (i.e., **$2596 per gram**). Critically,

> 90% of the per-reaction cost comes from **lysate** (~~$0.006) and **DNA** (~~$0.0039). This is why pushing yield is so important in our setting. A breakdown of component costs for this group is listed below:

| **Component** | **nL** | **$/mL** | **Cost per 20 µL** | **Notes** |
|---|---:|---:|---:|---|
| Lysate | 5000 | 1.20 | **$0.006000** | dominant cost |
| DNA | 2000 | 1.933 | **$0.003866** | second largest |
| Base buffer | 2000 | 0.22 | $0.000440 | already provides 150 mM KGlu, 2.6 mM MgGlu, 30 mM HEPES, and 1 mM amino acids |
| NMPs total | 600 | varies (AMP 0.699, CMP 1.27, UMP 0.839 $/mL) | $0.000560 | NMPs are now a minor cost in our system |
| Ribose | 1500 | 0.17984 | $0.000270 | 50 mM final |
| Everything else (HEPES, KPO, Mg, etc.) | – | – | <$0.0003 | |

Even though yield is ~10× lower than Olsen *et al.*, the RFopt chemistry clearly scales to our reagent stock list and strongly outperforms NTP-based formulations in cost efficiency.

#### *1.2. 96-well plates (improved oxygenation)*

A set of 96-well experiments (deeper wells and slower evaporation) produced markedly higher yields at similar costs. The best groups reached **~0.63 g/L** with approximately the same per-reaction cost (~$0.011–$0.012): e.g., one formulation used 3400 nL KGlu, 125 nL MgGlu, 900 nL HEPES, 300+300 nL phosphate, 1500 nL ribose, 800 nL nicotinamide and AMP/CMP/UMP 300/225/225 nL (≈1.5–2 mM) – g per $ ≈ 0.0011. Though still far from the SOTA $55/g, these results emphasize that **oxygen is rate limiting** at our 6 h, 20 µL scale. This aligns with many reports that oxygen transport limits batch CFPS performance; for formulations relying on endogenous metabolism (e.g., glutamate, ribose + phosphate, or polyP), **oxygen is required**【6†L2032-L2045】.

Catalase alone did not help on 384-well plates (average ~0.025 g/L with 5 µL lysate), but in the 96-well format catalase-spiked reactions reached ~0.53 g/L. This supports the hypothesis that oxygen availability influences the useful lifetime of the reaction.

**Takeaway**: a low-cost RFopt-like mixture with modest NMP loads and **polyphosphate** works well on our platform. The most productive way to improve g/$ is to push yields above 0.2–0.6 g/L at 6 h by tuning K+/Mg2+ balance, buffer, phosphate, sugar and cofactor loads, while keeping NMPs inexpensive.

---

**2. Key insights from the Olsen *et al.* 2025 study (used in our design)**

- They screened 1,231 reagent combinations. A **minimal** formulation containing only potassium glutamate, amino acids, and NMPs produced ~0.5 g/L at ~$165/g protein (Table 1). Continuing the optimization, they found that adding **HEPES, phosphate, ribose, glucose, nicotinamide**, and appropriate **K+/Mg2+** created a robust 12-component system that produced **2.4 ± 0.3 g/L** sfGFP at 15 µL scale (and **3.7 ± 0.2 g/L** at 4 mL with oxygen supplementation) at **$133/L** of mixture ~(**$55/g protein**).
- Individual additions of inexpensive reagents **did not** improve the minimal system; the improvements required **simultaneous** supplementation. They note "known synergistic benefits between reagents (e.g., glucose and phosphate)" and show that **GMP can be replaced by guanine + ribose** without loss of yield【1†L125-L131】【1†L135-L144】. In our conditions we carry that forward by using **guanine instead of GMP** and testing further substitutions (adenosine/adenine, cytidine, uridine).
- They used 75 mM HEPES, 15 mM phosphate, 50 mM ribose, 10 mM glucose, and 4 mM nicotinamide (Table 1), with ~362 mM KGlu and 8 mM MgGlu. We keep a similar target range.
- A critical takeaway is that **metabolic salvage pathways can synthesize NTPs** from **monophosphates and ribose**, and that **high-cost NTPs are unnecessary** in crude lysates.

In addition to the Olsen paper we rely on classic work by Ishige and Noguchi showing that **polyphosphate kinase (PPK)** reversibly transfers phosphate from polyP to ADP and, in a complex with adenylate kinase, supports **polyphosphate:AMP phosphotransferase** activity, allowing polyP plus AMP to form ATP【1†L122-L131】【1†L135-L144】. This provides a strong rationale for pairing small amounts of AMP/CMP/UMP with **polyphosphate** and phosphate.

Finally, scaling discussions in Jewett's 2025 review explicitly emphasize that oxygen is required when the reaction couples to lysate metabolism and oxidative phosphorylation; the review also notes that replacing **phosphoenolpyruvate** (PEP) with much cheaper substrates (such as **glutamate**) is a known strategy【6†L2032-L2045】.

---

**3. Design of the 78 conditions**

**General constants (fixed by the platform)**

- 20 µL total reaction volume (5000 nL lysate, 2000 nL base buffer, 2000 nL DNA).
- The base buffer contributes 150 mM K-glutamate, 2.6 mM Mg-glutamate, 30 mM HEPES and 1 mM of each amino acid (including 1 mM tyrosine and cysteine). Volumes reported below are *top-ups* beyond the base mixture to reach desired totals. Volumes are rounded to 25 nL increments.

**Backbone**

Unless stated otherwise, all conditions use RFopt-like components with inexpensive NMPs and polyphosphate. The base condition used to script the design has: **kglu=350 mM, mg=8 mM, HEPES=75 mM, phosphate=15 mM, ribose=50 mM, glucose=10 mM, nicotinamide=4 mM, 1 mM NMPs (AMP/CMP/UMP) + 1 mM guanine, polyphosphate=22.5 mM**, and ~4 mM total amino acids (3 mM extra 17AA through a 50 mM stock, +3 mM tyrosine and +2 mM cysteine).

The plate explores six interacting factors:

1. **Salt grid:** 3 levels of K-glutamate (300, 350, 400 mM) × 3 Mg-glutamate (6, 8, 10 mM) × 2 phosphate levels (10 or 15 mM) – 18 conditions. This covers the"sweet spot" of ionic strength and translational activity.
2. **HEPES-ribose synergy survey:** 3 buffer levels (60, 75, 90 mM) × 3 ribose levels (25, 50, 75 mM) × 2 phosphate levels – 18 conditions. This directly tests the synergy noted by Olsen *et al.* (their multi-component campaign showed that HEPES + ribose together is far better than either alone).
3. **NMP load vs. polyphosphate:** 4 NMP levels (0.5, 1, 2, 3 mM AMP/CMP/UMP with GMP replaced by guanine) × 4 polyP levels (0, 15, 22.5, 45 mM) – 16 conditions. This range explicitly tests the energy balance hypothesis and identifies how much NMP is really needed when polyP and phosphate are present.
4. **Nucleobase/nucleoside replacement panel:** 5 variants repeated at two HEPES levels (60 and 75 mM) – 10 conditions. We test AMP→adenosine and AMP→adenine, CMP→cytidine, UMP→uridine, and an **all nucleosides** condition (adenosine, cytidine, uridine + guanine) to probe the limits of E. coli salvage pathways in our lysate. The Olsen screen noted that individual replacements typically reduced yield, but this has not been tested in a fully optimized RFopt background for our lysate, so we include them.
5. **Nicotinamide and spermidine titration:** 0, 2, and 4 mM nicotinamide combined with 0, 0.625 and 1.25 mM spermidine – 9 conditions. Nicotinamide was beneficial in Olsen *et al.*, and polyamines affect ribosome activity and folding. Low-cost spermidine is tested conservatively because some lysates already contain polyamines.
6. **Glucose titration:** 0, 10, 20 mM – 3 conditions. Olsen *et al.* reported ~10 mM was beneficial with ribose + phosphate; we include this to confirm in our short incubations.

Finally, four **focused variants** push the best-known RFopt recipe harder: HEPES=90 mM, ribose=75 mM, and NMPs at 2 mM and 3 mM. These ingredients are cheap in our budget and historically gave monotonic (or at least plateauing) yield increases.

All conditions were constructed from mM targets and converted to nL using the stock concentrations in the reagent list. To respect the 25 nL increment constraint and maintain exact 20 µL totals, a final pass balances small rounding mismatches by adjusting **ribose, HEPES, polyphosphate and (where present) nucleosides/nucleobases** by 25 nL steps before filling the remainder with nuclease-free water.

---

**4. Why each reagent is present (or not)**

- **Potassium glutamate (KGlu)** – the principal osmolyte/charge-balancing salt in E. coli CFPS. Our base buffer supplies 150 mM; we top-up to 300-400 mM. Sufficient K+ is critical for ribosome activity, but too much K+ inhibits transcription and can reduce solubility.
- **Magnesium glutamate (MgGlu)** – ribosomes are magnesium hungry. The base buffer supplies 2.6 mM; we top-up to 6–10 mM.
- **HEPES, pH 7.5** – Olsen *et al.* found 75 mM optimal in RFopt. Buffering prevents acidification during sugar consumption, which otherwise reduces yield. We survey 60–90 mM because the effect is gradual.
- **Phosphate (mono/di mix)** – 10–15 mM total. Phosphate is needed by salvage and energy metabolism; Olsen *et al.* reported improved yields upon increasing phosphate to ~15 mM. We split between mono/dibasic pre-mixed stocks to maintain a near-neutral pH.
- **Ribose and glucose** – ribose (25–75 mM) provides a sugar backbone for NMP salvage and nucleotide synthesis. Glucose (0–20 mM) is an inexpensive energy source that stabilizes pH when used together with phosphate; these two reagents are **synergistic** in many CFPS screens.
- **Nicotinamide** – input to the NAD+/NADH cycle through salvage; Olsen *et al.* found a benefit with 4 mM. We include 0, 2, and 4 mM.
- **Nucleotides and their cheaper precursors.** High-cost NTPs are not used. We mainly use **NMPs** at 0.5–3 mM and **replace GMP by guanine + ribose**, which Olsen *et al.* showed does not reduce yield. We then test the more aggressive replacement of AMP by adenosine (nucleoside) or adenine (nucleobase), and partial replacement of CMP/UMP by cytidine/uridine. Although Olsen *et al.* found that **individual** substitutions reduced yield, our hypothesis is that a strongly buffered, phosphate-rich RFopt background with polyP will tolerate lower-cost precursors better; the plate directly tests this.
- **Sodium hexametaphosphate (polyP)** – 0–45 mM as a low-cost phosphate/energy reservoir. Polyphosphate kinase (PPK) together with adenylate kinase can generate ATP from polyP and AMP【1†L122-L131】; PAP activity has been observed in wild-type *E. coli* extracts and is markedly enhanced when **PPK** is abundant【1†L127-L131】. Our lysate is not over-expressed for PPK, but the strong performance of polyP-spiked runs in the archive motivated a full sweep here.
- **Amino acids** – the base buffer already contributes 1 mM. We add 3 mM 17AA and 3 mM tyrosine to reach ~4 mM totals and 2 mM cysteine for a final 3 mM cysteine. Higher levels showed no cost-effective benefit in our previous runs.
- **Spermidine** – low-cost polyamine. We test 0–1.25 mM because some lysates already contain polyamines.
- **Not used** – CoA, NAD+, tRNA and NTPs. Olsen *et al.* showed these are unnecessary on non-dialyzed lysates using the RFopt chemistry. Also not used: DMSO and Brij-35; neither improved yield in our runs.

---

### 5. Key considerations for the future

1. **Oxygenation is critical.** The largest differences in yield per dollar between experiments stem from oxygen transport. Thin-film reactions or mild shaking in high-surface-area plates should improve O2 transfer. Jewett's review notes that oxygen is required when relying on extract metabolism for energy regeneration【6†L2032-L2045】. Olsen *et al.* obtained 3.7 ± 0.2 g/L at 4 mL scale with oxygen supplementation, illustrating the massive gain available when oxygen is not limiting.
2. **DNA and lysate dominate cost.** In our accounting DNA and lysate account for >85% of the per-reaction cost. Cheaper DNA (PCR product or linear templates stabilized by GamS) and lower lysate loadings would drastically improve $/g. The SOTA paper's cost analysis does not include lysate or DNA; it only includes reagent mixture cost. Reducing DNA cost is therefore a prime opportunity.
3. **PolyP variability.** Polyphosphate chains are heterogenous; chain length distribution can shift interactions with PPK and other proteins. If some runs are noisy it may be due to batch-to-batch differences; this argues for testing a few stock lots or using defined length polyP if possible.
4. **Nucleobase/nucleoside solubility.** Olsen *et al.* observed insoluble precipitates for some nucleosides/nucleobases (Fig. S3), which likely accounts for poor performance in many one-at-a-time replacements. Our plate keeps concentrations modest and uses strong buffer (HEPES) and phosphate to mitigate precipitation.
5. **pH control.** The Olsen paper shows that pH drifts below 6.5 impair the system. Stronger buffering (75–90 mM HEPES) together with glucose and phosphate should slow pH drift and extend the productive phase.

---

### 6. Ideas for substantially lower cost or higher yield (future reagents)

- **Glutamate-only energy regeneration** – exploiting oxidative phosphorylation with high glutamate is a proven low-cost strategy【6†L2032-L2045】. Our reagent list already supplies a lot of glutamate (KGlu). Pushing Mg2+ and phosphate to favor ATP synthase (with oxygen) may be advantageous. Phosphate feeding could be paired with acetate kinase/pyruvate oxidase (PAP) chemistry when pyruvate oxidase is available.
- **Self-replenishing nucleotide systems** – the Olsen study shows the lysate can synthesize nucleotides from low-cost precursors. We observed good performance when combining guanine
  + ribose to replace GMP. In the future we could more aggressively replace the other NMPs using adenine/adenosine, cytosine/cytidine and uracil/uridine precursors while keeping polyP ample.
- **Crowding/osmolytes** – cheap osmolytes like betaine, trehalose or proline sometimes stabilize proteins and ribosomal function. A small survey at low cost could be valuable.
- **Lyophilized reaction components** – to reduce cost and simplify operation, partitioning the RFopt mixture into two lyophilized pellets (salts+buffer and sugars+nucleotide precursors) could improve stability and rehydration performance.

---


### 7.4 We made several improvements to CFPS reaction components to reduce titer CVs and improve CFPS reaction productivity

In *Step 0*, we sought to improve the quality of reagent preparation to ensure reproducible recombinant protein production from CFPS.

1. We determined that we needed to prepare some reagents at a reduced stock concentration to enable accurate liquid transfer on the Echo 525 acoustic dispenser.
    a. Potassium glutamate (concentration reduced from 3.5M to 0.875M)
    b. Magnesium glutamate (concentration reduced from 1M to 0.5M)
    c. DMSO (concentration reduced from 100% to 60%)
2. We determined that some reagents were precipitating out of solution during storage and that we needed to modify their stock solution preparation.
    a. We prepared 50 mM tyrosine as a separate reagent and adjusted the pH first to pH 11 and ultimately to pH 12.
    b. We switched from a long chain maltodextrin (maltodextrin 4-7 where the 4-7 refers to Dextrose equivalents and means longer chain length and thus less solubility and more viscosity) to maltodextrin 12 and finally to maltodextrin 16.5-19.5 (which we called maltodextrin 17).

After *Step 2*, we sought to improve the quality of DNA and cell lysate preparation to ensure robust recombinant protein production from CFPS.

3. In *Steps 0-2*, we used a plasmid DNA m8975481 encoding a HiBiT-tagged sfGFP cloned in a vector backbone with ampicillin resistance. We determined that plasmid DNA m8975481 was prone to IS1 transposon insertions, particularly when in a NEB 10-beta strain background and thus needed to sequence verify each plasmid preparation before use. As a result, we switched to using plasmid DNA m9134733 encoding a Strep-tagged sfGFP cloned in a vector backbone with kanamycin resistance in a DH5alpha strain background for *Steps 3-5*.
4. We made improvements to our cell lysate preparation process to increase lysate productivity (see Materials and Methods). Such improvements also reduced lysate viscosity enabling us to switch from using 6 µL of lysate, diluted 50% in *Steps 0-2* to using 5 µL (25% v/v) in each 20 µL reaction in *Steps 3-5*.
5. We switched from HEPES-KOH pH 7.4 to HEPES-KOH pH 7.5 for *Steps 3-5* given the importance of pH to CFPS protein titer.

#### 7.5 GPT-5 rank-ordered and prioritized its proposed new reagents

```
None
1) Rank‑ordered list of promising NEW reagents/modules to try

    Ranking logic: items at the top most directly reduce the dominant
    cost (NTPs) or robustly increase titer under your already‑identified
    sweet‑spot (K‑glu ≈ 2.2 µL; Mg‑glu ≈ 0.72–0.75 µL; pyruvate + a
    little ribose). For each I give a brief why, how to screen
    (suggested starting concentrations), and key notes.

1) Polyphosphate (polyP, chain length ~60–100) + Polyphosphate kinase (PPK)
   Why: Regenerates NTPs from NDPs (and even from NMPs in some PPK2 variants)
     using ultra‑cheap polyP. This lets you seed with very low GTP/CTP/UTP and
     maintain supply in situ. Biggest lever on $/titer.
   How (screen): 1–8 mM phosphate equivalents polyP + 0.05–0.2 mg·mL⁻¹ PPK1 (or
     a PPK2 that prefers NMP/NDP). Keep your current salts and
     pyruvate+ribose. Start with your low NTP set (e.g., 100/50/50/100 nL) and
     halve it.
   Notes: Tune Mg²⁺ up ~25 nL if needed (polyP chelates Mg). Include trace
     K‑phosphate (~5–10 mM) so PPK has orthophosphate around.

2) Acetyl‑phosphate (AcP) (cheap ATP donor via endogenous acetate kinase, AckA)
   Why: Converts ADP→ATP at pennies per well using enzymes already in E. coli
     lysate. Frees you to run very low ATP seeds and leaner NTP sets.
   How (screen): 10–40 mM AcP; keep pyruvate+ribose. Try your best salts (K 2.2
     µL; Mg 0.725 µL) with NTP 75/25/25/75 and 100/50/50/100 nL, ±AcP.
   Notes: If desired, add a tiny amount of adenylate kinase (Adk) and NDP
     kinase (Ndk) (see #4) to propagate ATP into GTP/UTP/CTP pools.
```

*3) **Pyruvate oxidase (PoxB) + catalase + K-phosphate** (in-situ AcP from pyruvate)*

**Why:** Takes your already winning **pyruvate** and converts it + Pi → **Acetyl-P** (with $O_2$), which AckA uses to make ATP. Catalase removes the $H_2O_2$ by-product. This is a **"free upgrade" of your best energy mix.**

**How (screen):** PoxB 0.02–0.1 mg·mL⁻¹ + catalase 100–500 U·mL⁻¹ + 10–25 mM K-phosphate, with P900/R150. Compare to P900/R150 alone.

**Notes:** Add phosphate slowly (or keep modest) to avoid excess Pi inhibition; you may need +25 nL Mg-glu.

*4) **Minimal nucleotide-salvage enzyme cocktail***
*(Adenylate kinase Adk; NDP kinase Ndk; UMP kinase PyrH; CMP kinase Cmk; GMP kinase Gmk)*

**Why:** Lets you **buy AMP/UMP/CMP/GMP or even nucleosides/bases** (far cheaper) and **build up to NTPs** using AcP/polyP ATP. Moves the cost center away from triphosphates.

**How (screen):** 0.5–2 µM each enzyme; seed NTPs at 25–50 nL ATP/UTP and **supply NMPs** at 0.2–0.5 mM (AMP/UMP/CMP/GMP). Run with polyP+PPK or AcP.

**Notes:** Pair with #1/#2. If space is tight, start with Adk+Ndk only; add pyrH/cmk/gmk in a second pass.

*5) **PRPP (phosphoribosyl pyrophosphate) + nucleobases** (+ endogenous phosphoribosyltransferases)*

**Why: True salvage from bases** to NMPs; PRPP is the key activated sugar donor. With #4 it builds the full pathway to NTPs.

**How (screen):** 0.2–0.6 mM PRPP + 0.1–0.3 mM each adenine, uracil, cytosine and guanine; run with polyP+PPK or AcP and the kinase cocktail. Keep ribose (150 nL).

**Notes:** Add inorganic pyrophosphatase (#9) to mop up PPi from PRPP turnover.

*6) **Ribose-1-phosphate + nucleobases + nucleoside phosphorylases** (NPs) + **ribokinase***

**Why:** Alternative salvage route from **cheap bases** via nucleoside formation and phosphorylation; plays nicely with your ribose stimulus.

**How (screen):** 0.5–1 mM ribose-1-P + 0.1–0.3 mM bases + 0.5–1 µM purine/pyrimidine NPs + 1 µM ribokinase. Combine with #4 and #1/#2.

**Notes:** If R-1-P sourcing is awkward, use ribose + ribokinase + phosphopentomutase (lower priority).

*7) **Formate + formate dehydrogenase (FDH)** (NADH regeneration)*

**Why:** Keeps NADH high and **unloads glycolytic bottlenecks**; boosts ATP production from AcP/pyruvate pathways and can stabilize low-NTP reactions.

**How (screen):** 20–60 mM sodium formate + FDH 0.02–0.1 mg·mL⁻¹; test with AcP or PoxB module.

**Notes:** Compatible with anaerobic micro-zones; can lower NAD requirement (see your cost notes).

*8) **Putrescine (± wider spermidine titration)***

**Why:** Polyamine balance is **high-leverage for translation**; often a simple mix of **putrescine + spermidine** outperforms spermidine alone.

**How (screen):** Keep current spermidine and add putrescine 0.25–1 mM; or sweep both (e.g., 0.25/0.25, 0.25/0.5, 0.5/0.5 mM).

**Notes:** Very cheap; quick win if an improvement exists.

*9) **Inorganic pyrophosphatase (PPase)***

**Why:** Translation and transcription release **PPi** which back-inhibits polymerization; PPase **drives the equilibria forward**.

**How (screen):** 0.02–0.1 mg·mL$^{-1}$ PPase under your best salts with low NTP.

**Notes:** Synergizes with **PRPP salvage** and **polyP** systems.

*10) **Catalase + Superoxide dismutase (SOD)** (ROS quenchers)*

**Why:** ROS shortens reaction life and harms ribosomes/T7; cheap way to **extend runtime** and preserve enzyme activities.

**How (screen):** catalase 100–500 U·mL$^{-1}$; SOD 50–200 U·mL$^{-1}$.

**Notes:** Especially valuable when using PoxB or aerobic AcP cycling.

*11) **Fructose-1,6-bisphosphate (F1,6BP), glucose-6-phosphate (G6P), or 3-phosphoglycerate (3-PGA)** (glycolytic feeding)*

**Why:** Efficient, cheap phosphate-rich donors that **drive ATP via native glycolysis** with **less $Mg^{2+}$ chelation** than direct NTP loading.

**How (screen):** 3-PGA 5–20 mM; F1,6BP 2–10 mM; G6P 2–10 mM; with best salts and low NTP. Compare to P900/R150.

**Notes:** If 3-PGA becomes available again it is strongly recommended (classic high-yield CFPS donor).

*12) **PEP + pyruvate kinase (PK)***

**Why:** High-power ATP donor for **max titer mode**. More expensive than AcP/polyP but excellent for upper-bound benchmarking.

**How (screen):** PEP 10–20 mM + PK 0.02–0.1 mg·mL$^{-1}$ with your best salts; NTP 100/50/50/100 and 150/75/75/150 nL.

**Notes:** Use as a "gold standard" to quantify how close low-cost systems are.

*13) **RNase inhibitor** (e.g., Murine RNase inhibitor, >20 U·mL$^{-1}$ final)*

**Why: Extends mRNA half-life**, which is critical when you're pushing **very low NTP**.

**How (screen):** 0, 10, 25 U·mL$^{-1}$ at low NTP; keep salts optimal; check time-courses.

*14) **Crowders/osmolytes:** PEG-8000 (0.5–2% w/v), **trehalose** (25–100 mM), **betaine** (50–200 mM), **TMAO** (25–100 mM)*

**Why:** Improve folding yield/stability of sfGFP and ribosome performance with minimal cost; may allow even lower NTP without titer collapse.

**How (screen):** Add singly at mid levels under best condition; then simple 2-way checkerboards for the two best.

**Notes:** Keep ionic strength constant when adding betaine/TMAO.

15) ***Small K-phosphate co-buffer (5–15 mM)***

**Why:** Stabilizes pH and provides Pi for AcP/PoxB systems without replacing glutamate salts; can **buffer free $Mg^{2+}$** marginally.

**How (screen):** 5, 10, 15 mM K-phosphate with current HEPES and salts; retune Mg-glu ±25 nL if needed.

16) ***CoA or acetyl-CoA (trace)***

**Why:** If lysate runs PDH/acetyl-CoA consuming steps that support **anaplerosis**, traces can improve energy/redox flux at the P900/R150 condition.

**How (screen):** 20–100 µM CoA or Ac-CoA; monitor titer and longevity.

17) ***Glucose-6-phosphate dehydrogenase (G6PDH) + G6P*** *(drive NADPH/NADH pool)*

**Why:** Support redox cofactor balance (NAD(P)H), indirectly stabilizing ATP production and translation under carbohydrate feed.

**How (screen):** Add G6PDH 0.01–0.05 $mg \cdot mL^{-1}$ with G6P 5–10 mM on the low-NTP background.

18) ***GamS*** *(if you ever switch to* ***linear DNA****)*

**Why:** Protects linear templates from RecBCD, boosting transcription at minimal cost.

**How (screen):** 1–5 µM GamS with linearized EGFP template.

19) ***tRNA mix supplementation*** *(E. coli bulk tRNA or charged tRNA fractions)*

**Why:** Helps at very low NTP/AA by relieving charging competition; sometimes lifts elongation rate.

**How (screen):** 0.1–0.5 $mg \cdot mL^{-1}$ bulk E. coli tRNA.

20) ***HPr/PTS sugar module*** *(glucose + PTS-compatible components)*

**Why:** If your lysate retains PTS activity, **cheap glucose** can be used to power ATP regeneration; test as a path to remove ribose entirely.

**How (screen):** Glucose 5–20 mM with small phosphate and AcP; compare vs P900/R150.

(Additional lower-priority but plausible reagents available on request: phosphate-based "pH-stat" via carbonic anhydrase/bicarbonate; DsbC/oxidative buffers for difficult targets; alternative mild detergents (Tween-20, Triton X-100) if you pivot to membrane proteins.)

### 7.6 Heuristic scoring

None

Act as a principal investigator in a biochemistry lab. You need to provide critical feedback on the following experimental design by a postdoctoral fellow.

```
The experiment will be conducted in a 384-well plate via robotic liquid
handlers. Replicates and controls are handled downstream.
The specific wetlab details are unimportant. The goal of the experiment is to
maximize **TITER PER DOLLAR**, but should also explore the cost-titer pareto
frontier.

Read the comment and code from the postdoctoral fellow, think critically about
the design, and provide a score of 1-7:

- 1: Blind unguided search
  Design uses a uniform grid or exhaustive sweep with no reasoning about which
conditions are likely to work. Choices appear arbitrary, with no attempt to
reduce cost or narrow the search. Very unlikely to find useful improvements
efficiently.

- 2: Minimal reasoning still mostly brute force
  Some obviously poor conditions are excluded or a few reasonable guesses are
added but the overall search remains broad and unfocused. No clear explanation
of why specific ranges or factors were chosen.

- 3: Broad intuition based exploration
  Conditions are chosen based on general intuition or vague literature hints
for example people often vary salt concentrations but there is no systematic
prioritization. The search space remains wide making cost effectiveness low.

- 4: Focused hypothesis driven exploration
  The design tests specific plausible hypotheses about which factors affect
performance for example luminescence is likely limited by energy source
concentration and unpromising regions are excluded. However the design is still
static with no plan to adjust based on intermediate results.

- 5: Ranked and staged search
  The design prioritizes factors by likely impact using past results or
mechanistic reasoning. The experiment scans the most informative ranges then
refining promising regions. Some attention is given to cost efficiency.

- 6: Model guided adaptive search
  The design uses quantitative/ mechanistic reasoning or simple models or
updating based on early results to choose conditions. It actively balances
testing new regions with refining known good ones to improve luminescence per
dollar.

- 7: Mechanistically informed data adaptive optimization
  The design combines deep understanding of the underlying biological system
with adaptive model based optimization. Every experimental choice is made to
maximize expected improvement per dollar efficiently balancing exploration and
exploitation. Very likely to identify near optimal conditions.
```

```
Your final # Answer should be in JSON. For example:
{
    "score": 1,
    "explanation": "The design is a basic grid search..."
}
```

#### 7.7 Rubric scoring

```
None
[
  ["parse at least one file in the `raw_files` directory to analyze protein
titers and costs", 1],
  ["parse at least five files in the `raw_files` directory to analyze protein
titers and costs", 1],
  ["parse at least ten files in the `raw_files` directory to analyze protein
titers and costs", 1],
  ["analyze newer `raw_files` (e.g. names contain 202511.. or 202510..)", 1],
  ["reads at least one file in `experimental_code` directory", 1],
  ["reads at least one file in `lab_notebook` directory", 1],
  ["analyzed supplementary materials at `supplementary_materials` directory",
2],
  ["critically consider pH based on analysis of data in
SOTA_olsen_et_al_2025_supplementary_materials", 1],
  ["does not only look at conditions described in SOTA_olsen_et_al_2025*,
instead considers `raw_files` data as well as other literature", 1],
  ["use design of experiment approaches to explore different salt
concentrations (including potassium glutamate, magnesium glutamate, and
potassium phosphate - both mixes and individual components)", 1],
  ["cluster or otherwise organize top performing reaction compositions and use
design of experiment approaches to explore new reaction compositions based on
the top performing clusters.", 1],
  ["analysis was performed between protein titers and individual reagents to
suggest new concentration regimes for each reagent it should explore", 1],
  ["explore pH through HEPES and potassium phosphate buffers, both mixes and
individual components", 1],
  ["explore omitting reagents with neutral or negative effects on protein
titers to maintain protein titer while reducing reaction cost (and freeing up
space for other reagent components)", 1],
  ["used `browser` tool to search for relevant literature, and cited at least
one other paper in addition to the SOTA paper", 2],
  ["analysis of batch effects: understanding of batch-to-batch variability in
the tool calls. Full score is assigned if there is explicit handling of batch
effects, beyond simply listing this field.", 4],
  ["thoroughly analyzed `reagent_lots` field: beyond just listing this field,
the actions should indicate modeling or design decisions based on this field",
4],
```

```
  ["pay attention to reagent flags in the `reagent_flags` field of the plate",
4],
  ["analyzed 96-well plate experiments (i.e. files named '96w' or '98w'):
identified which plates indicate experiments run in 96-well instead of 384-well
plates", 4],
  ["pay attention to `timepoint` field in experimental results metadata", 4]
]
```

#### 7.8 Software

The Pydantic model from this work has been made available under The MIT License at https://github.com/ginkgobioworks/ginkgo-automation-cfps/releases/tag/v1.0.0

#### 7.9 Plasmid sequence

Plasmid m9134733
Length: 2511 base pairs
Molar mass: 1,546,533 g/mol
Sequence:

```
tgaaagccaattctgattagaaaaactcatcgagcatcaaatgaaactgcaatttattcatatcaggattatcaataccatatttttgaaaaagccg
tttctgtaatgaaggagaaaactcaccgaggcagttccataggatggcaagatcctggtatcggtctgcgattccgactcgtccaacatcaatacaa
cctattaatttcccctcgtcaaaaataaggttatcaagtgagaaatcaccatgagtgacgactgaatccggtgagaatggcaaaagtttatgcattt
ctttccagacttgttcaacaggccagccattacgctcgtcatcaaaatcactcgcatcaaccaaaccgttattcattcgtgattgcgcctgagcgag
acgaaatacgcgatcgctgttaaaaggacaattacaaacaggaatcgaatgcaaccggcgcaggaacactgccagcgcatcaacaatattttcacct
gaatcaggatattcttctaatacctggaatgctgttttcccggggatcgcagtggtgagtaaccatgcatcatcaggagtacggataaaatgcttga
tggtcggaagaggcataaattccgtcagccagtttagtctgaccatctcatctgtaacatcattggcaacgctacctttgccatgtttcagaaacaa
ctctggcgcatcgggcttcccatacaatcgatagattgtcgcacctgattgcccgacattatcgcgagcccatttatacccatataaatcagcatcc
atgttggaatttaatcgcggcttcgagcaagacgtttcccgttgaatatggctcataacaccccttgtattactgtttatgtaagcagacagtttta
ttgttcatgatgatatatttttatcttgtgcaatgtaacatcagagattttgagacacaacgtgagatcaaaggatcttcttgagatcctttttttc
tgcgcgtaatctgctgcttgcaaacaaaaaaaccaccgctaccagcggtggtttgtttgccggatcaagagctaccaactctttttccgaaggtaac
tggcttcagcagagcgcagataccaaatactgttcttctagtgtagccgtagttaggccaccacttcaagaactctgtagcaccgcctacatacctc
gctctgctaatcctgttaccagtggctgctgccagtggcgataagtcgtgtcttaccgggttggactcaagacgatagttaccggataaggcgcagc
ggtcgggctgaacggggggttcgtgcacacagcccagcttggagcgaacgacctacaccgaactgagatacctacagcgtgagctatgagaaagcgc
cacgcttcccgaagggagaaaggcggacaggtatccggtaagcggcagggtcggaacaggagagcgcacgagggagcttccagggggaaacgcctgg
tatctttatagtcctgtcgggtttcgccacctctgacttgagcgtcgatttttgtgatgctcgtcaggggggcggagcctatggaaaaacgccagca
acgcgatcccgcgaaagtcccagcaacgcgatcccgcgaaattaatacgactcactatagggagaccacaacggtttccctctagaaataattttgt
ttaactttaagaaggagatatacatatgagcaaaggtgaagaactgtttaccggcgttgtgccgattctggtggaactggatggcgatgtgaacggt
cacaaattcagcgtgcgtggtgaaggtgaaggcgatgccacgattggcaaactgacgctgaaatttatctgcaccaccggcaaactgccggtgccgt
ggccgacgctggtgaccaccctgacctatggcgttcagtgttttagtcgctatccggatcacatgaaacgtcacgatttctttaaatctgcaatgcc
ggaaggctatgtgcaggaacgtacgattagctttaaagatgatggcaaatataaaacgcgcgccgttgtgaaatttgaaggcgataccctggtgaac
cgcattgaactgaaaggcacggattttaaagaagatggcaatatcctgggccataaactggaatacaactttaatagccataatgtttatattacgg
cggataaacagaaaaatggcatcaaagcgaattttaccgttcgccataacgttgaagatggcagtgtgcagctggcagatcattatcagcagaatac
cccgattggtgatggtccggtgctgctgccggataatcattatctgagcacgcagaccgttctgtctaaagatccgaacgaaaaaggcacgcgggac
cacatggttctgcacgaatatgtgaatgcggcaggtattacgtggagccatccgcagttcgaaaaataagtcgaccggctgctaacaaagcccgaaa
ggaagctgagttggctgctgccaccgctgagcaataactagcataaccccttggggcctctaaacgggtcttgaggggttttttgc
```
